# Copy-number intratumor heterogeneity contributes to predict relapse in chemotherapy-naive stage II colon cancer

**DOI:** 10.1101/2021.04.16.440177

**Authors:** Sara Lahoz, Iván Archilla, Elena Asensio, Eva Hernández-Illán, Queralt Ferrer, Sandra López-Prades, Ferran Nadeu, Javier del Rey, Rebeca Sanz-Pamplona, Juan José Lozano, Antoni Castells, Miriam Cuatrecasas, Jordi Camps

## Abstract

Optimal selection of high-risk patients with stage II colon cancer is crucial to ensure clinical benefit of adjuvant chemotherapy after surgery. Here, we investigate the prognostic and predictive value of genomic intratumor heterogeneity and aneuploidy for disease recurrence. We combined SNP arrays, targeted next-generation sequencing, fluorescence in situ hybridization and inmunohistochemistry on a retrospective cohort of 84 untreated stage II colon cancer patients. We assessed the subclonality of copy-number alterations (CNAs) and mutations, CD8+ lymphocyte infiltration and their association with time to recurrence (TTR). Prognostic factors were included in machine learning analysis to evaluate their ability to predict individual relapse risk. Tumors from recurrent patients (N = 38) exhibited greater proportion of CNAs compared with non-recurrent (N = 46) (mean 31.3% vs. 23%, respectively; *P* = 0.014), which was confirmed in an independent cohort. Candidate chromosome-specific aberrations included the gain of the chromosome arm 13q (*P* = 0.02; HR, 2.67) and loss of heterozygosity at 17q22-q24.3 (*P* = 0.05; HR, 2.69), both associated with shorter TTR. CNA load positively correlated with intratumor heterogeneity (*R* = 0.52; *P* < 0.0001), indicating ongoing chromosomal instability. Consistently, subclonal copy-number heterogeneity was associated with elevated risk of relapse (*P* = 0.028; HR, 2.20), which we did not observe for subclonal mutations. The clinico-genomic model rated an area under the curve of 0.83, achieving a 10% incremental gain compared to clinicopathological markers. In conclusion, tumor aneuploidy and copy-number heterogeneity were predictive of a poor outcome in early-stage colon cancer, and improved discriminative performance in comparison to clinicopathological data.

## INTRODUCTION

Early-stage colon cancer holds a major therapeutic challenge due to the lack of strong biomarkers to predict disease recurrence [1]. Around 10 to 15% of patients diagnosed with stage II colon cancer relapse within the five years after curative intended surgery, which compromises survival. Although extensive biomarker-driven research has succeeded in identifying risk predictors for tumor dissemination, selection of patients for adjuvant chemotherapy is still controversial in stage II colon cancer [2].

To date, the most relevant clinical and pathological factors for discriminating high-risk stage II individuals are bowel perforation or obstruction, tumor size and high histological grade, lymphovascular or perineural invasion, and serosal invasion, although most of them have a modest individual effect on recurrence risk [3]. Microsatellite instability (MSI) is one of the most solid molecular markers with clinical utility in non-advanced colon cancer, and along with *BRAF*^V600E^ mutation, both are able to define a subgroup of stage II colon cancer patients (~6%) with improved survival when solely treated with surgery [4]. Tumor budding and the presence of poorly differentiated clusters are pathological-based markers that also contribute to the prognostication of these patients [5]. Moreover, Immunoscore is a robust immunohistochemistry-based prognostic index for quantifying tumor infiltrating CD8+ T cells that has been thoroughly validated in stages II-III colorectal cancer (CRC) [6,7]. Notwithstanding, recommendations in clinical guidelines for risk stratification still rely on under-sensitive clinical and histopathological criteria.

Genomic intratumor heterogeneity has been occasionally appointed as a prognostic predictor in solid malignancies [8,9] due to its putative ability of prompting somatic evolutionary processes that can drive cancer progression [10] and therapeutic failure [11,12]. Much of the genetic heterogeneity observed in solid tumors is triggered by chromosomal instability (CIN) [13], resulting in a pervasive expansion of chromosomal aneuploidies and mutational events that increases tumor subclonality and accelerates genome evolution [14,15]. Such instability could be responsible of fueling the tempo-spatial spread of relapse-proficient subclones that might compound the eventual metastatic seed [16,17], potentially explaining mechanisms of disease recurrence, such as those described in chronic lymphocytic leukemia [18]. In this sense, extensive copynumber subclonality has been associated with an increased risk of recurrence or death in non-small cell lung cancer [19] as well as in hepatocellular carcinoma [20]. Furthermore, ongoing CIN enables cancer cells to rapidly assemble complex karyotypes with inflated burdens of copy-number alterations (CNAs) [21], conferring the tumor a selective advantage towards a more aggressive phenotype [22,23]. Indeed, the tumor CNA load has been posed as a biomarker of reduced survival in prostate cancer [24] and in metastatic CRC was also predictive of a diminished response to bevacizumab combinatorial therapy [25]. Nevertheless, genomic complexity and intratumor heterogeneity are yet to materialize as clinically useful tools to monitor cancer progress for therapeutic decision-making.

Here, we perform a comprehensive integration of somatic CNAs and mutational events, their subclonal status, along with the CD8+ immune infiltration and other clinicopathological features in stage II colon cancer. By applying machine learningbased modeling using data from single nucleotide polymorphism (SNP) arrays, nextgeneration sequencing (NGS), fluorescence in situ hybridization (FISH) and immunohistochemistry, we suggest that genomic intratumor heterogeneity and aneuploidy are predictive determinants of tumor dissemination.

## MATERIALS AND METHODS

### Cohorts of patients and tumor samples

A total of 84 stage II colon adenocarcinomas (pT3-4N0M0) provided by the Hospital Clínic of Barcelona/IDIBAPS Tumor Biobank were retrospectively analyzed for this study. Patients were diagnosed between 2005 and 2016, and none of them received adjuvant chemotherapy after surgical resection of the primary tumor. Among them, 38 patients (45%) developed disease recurrence and 46 (55%) did not show recurrence after a median follow-up time of 7.6 years (range from <1 to 14.5 years). The primary endpoint was time to recurrence (TTR), defined as the time from surgery of the primary tumor to disease recurrence, where death without recurrence were censored at the time of death. This study was approved by the institutional ethics committee from Hospital Clínic of Barcelona (register HCB/2018/0174), and all patients signed an informed consent in accordance with the Declaration of Helsinki. For analytical validation purposes, an independent cohort of 99 untreated stage II microsatellite-stable (MSS) colon cancer patients (Colonomics) was included (EGAD00010001253) [26].

### SNP-arrays and copy-number analysis

Copy-number and loss-of-heterozygosity (LOH) profiling of tumors was performed using genome-wide Affymetrix OncoScan SNP-arrays on unmatched tumor specimens. All resulting CEL files were loaded onto Nexus Copy Number software version 9.0 (BioDiscovery, El Segundo, CA, USA) for data analysis and visualized using CNApp [27]. To generate whole-genome copy-number and allele-frequency calls, raw SNP-array fluorescence intensity data were processed using SNP-FASST2 segmentation algorithm from Nexus Biodiscovery software, achieving an estimate of the mean log2-ratio and B-allele frequency (BAF) values for each genome region. In-house resegmentation parameters included minimum segment length of 2Mb as well as joining of adjacent segments when separated <2.5Mb and having a log2-ratio/BAF difference <|0.1|. The CNApp tool [27] was employed to generate whole-genome plots displaying the frequency of copy-number alteration (CNA) accumulated in each chromosome region. For a chromosome region to be considered as carrying a CN gain or loss, log2-ratio cut-offs were set on 0.18 and −0.18, respectively; and to be considered to have a loss of heterozygosity (LOH), BAF threshold was set on 0.3. Chromosomal regions showing a non-altered log2-ratio and BAF <0.3 were classified as copy-number neutral LOH (CNN-LOH). Re-centralization of the zero copy-number line was performed manually in Nexus according to each chromosome BAF profile to re-adjust diploid regions onto the zero log2-ratio level. Later for each tumor, the proportion of the genome that was aberrant, or CNA load, was inferred from dividing the mean-length of segments with a log2-ratio ≥|0.18| by the summatory of lengths from all segments in that tumor genome.

Major and minor allele numbers of each segment were inferred utilizing ASCAT R package version 2.5 [28]. The proportion of tumor cells, or cancer cell fraction (CCF), carrying each CNA was estimated from ASCAT segmented data as a function of the B-allele frequency and the average major/minor numbers at each segment, adjusted by sample purity, using the following equation:

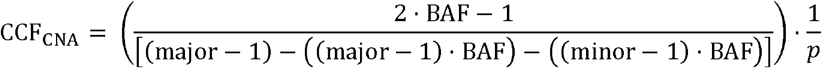

where BAF corresponds to the major frequency (>0.5) of the B-allele, *major* to the number of copies of the B-allele, *minor* to the number of copies of the A-allele, and *p* to the sample purity (0 to 1) obtained from ASCAT. Segments with CNA and copynumber neutral-LOH (CNN-LOH) were considered subclonal when their CCF was <85% and clonal otherwise [29], although more stringent CCF thresholds were also tested.

### Targeted next-generation sequencing and mutation analysis

The full-coding region of 48 CRC-related genes (Supplementary Table S1) was sequenced using a MiSeq platform (Illumina, San Diego, CA, USA) on a subset of 44 unpaired tumor samples. Raw sequencing reads on FASTQ files were submitted to base quality control and mapped to human reference genome GRCh37/hg19 utilizing the MiSeq Reporter software (MSR, v2.6, Illumina). Variant calling was executed using two algorithms: MuTect2 (v2.2) from the GATK platform (Broad Institute, Cambridge, MA, USA) and the Illumina Somatic Variant Caller (integrated in the MiSeq Reporter software, v2.6). Variants on VCF files were annotated with SnpEff (v4.3) plus dbNSFP (v2.9.3). Low quality calls were discarded to minimize false positive events: mapping quality below 30, TLOD <9 and read depth below 15x. Synonymous and germline variants were removed, along with polymorphisms having a worldwide population frequency above 1%, as reported in databases ExAC and 1000 Genomes Project. In addition to meeting the previous criteria, variants were manually curated and prioritized for final analysis only if: (i) they appeared in COSMIC and/or ClinVar databases as pathogenic, and/or (ii) were predicted as damaging events by at least 3 of 5 pathogenicity prediction scores (SIFT, PolyPhen2-HumVar, PolyPhen2-HumDiv, MutationTaster and MutationAssessor).

Given a mutation, the variant allele frequency (VAF) was converted into a CCF_mut_ using next equation [30], where the VAF is adjusted by the copy-number at the mutated locus and by sample purity (obtained from ASCAT):

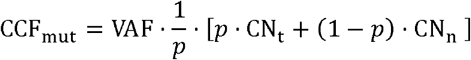

where *p* corresponds to the sample purity (0 to 1), CN_t_ to the tumor locus copy-number of each mutated coordinate, and CN_n_ to the normal locus copy-number (being assumed to be diploid, 2 copies, in autosomic chromosomes). Mutation events appearing in ≥85% of cancer cells were considered clonal, and subclonal otherwise.

### Predictive modeling by machine learning

Multivariable classification analyses were computed integrating all master co-variables to predict the probability of experiencing relapse for each patient. The gradient boosting machine algorithm was employed to classify patients. To predict missing values, multifold imputations were performed using *mice* R package. Numerical data were scaled and re-centered resulting in a joint distribution of mean 0 and standard deviation of 1. In order to minimize the overfitting bias, samples were randomly split into iterative training and testing datasets (in a 3:1 ratio). All fitted training models were cross-validated using a 4-fold procedure, and the outcome parameters used to predict their respective testing datasets using the *caret* R package. Several classifier algorithms were tested for the model generation: gradient boosting machine, random forest, LASSO, logistic regression, support vector machine (SVM) with linear and polynomial kernels, k-nearest neighbor (KNN) and linear discriminant analysis (LDA). To evaluate the predictive performance of the binary classifier: the sensitivity, specificity, and positive and negative predictive values (PPV and NPV, respectively), and associated 95% confidence intervals, were inferred using *crossval* R library. Diagnostic ability of the model was assessed by means of its area under the curve (AUC) on a receiver operating characteristic (ROC) curve with *pROC* package. To check whether the incremental gain on the AUC was significant between two different biomarker models, DeLong’s method on *pROC* package was used.

### Data availability

Additional information related to Material and Methods, including tissue microarrays, FISH, immunohistochemistry, prognostic modeling and statistical analysis is provided in Supplementary Methods. SNP-array and targeted next-generation sequencing data have been deposited in the National Center for Biotechnology Information (NCBI) Gene Expression Omnibus (GSE172191).

## RESULTS

### Clinicopathological characteristics and their prognostic value in the study population

Clinical and tumor-related histopathological characteristics are annotated in Table 1. Median age at diagnosis was 74 years old (range 55 – 91) and 54% of individuals were male. Most tumors were pT3 stage (73%) and had low histological grade (88%). Lymphovascular or perineural invasion and an invasive infiltrating margin were significantly more frequent in patients with disease relapse (*P* = 0.016 and 0.0008, respectively). Recurrent tumors showed remarkably lower amounts of tumor infiltrating CD8+ lymphocytes compared to non-recurrent (*P* = 0.0003) (Supplementary Figure S1A). High tumor budding and poorly differentiated clusters were also associated with disease relapse (*P* = 0.019 and 0.008, respectively).

**Table 1.** Patient and tumor-related clinicopathological characteristics.

| Variable | Strata | All (N=84) |  | Recurrent patients (N=38) |  | Non-Recurrent patients (N=46) |  | P value |
| --- | --- | --- | --- | --- | --- | --- | --- | --- |
|  |  | N | % | N | % | N | % |  |
| Age | Median, years (range) | 77 (55 – 91) |  | 79 (55 – 87) |  | 76 (55 – 91) |  | 0.24 |
| Sex | Male | 45 | 53.57% | 23 | 60.53% | 22 | 47.83% | 0.35 |
|  | Female | 39 | 46.43% | 15 | 39.47% | 24 | 52.17% |  |
| pT grade | T3 | 61 | 72.62% | 26 | 68.42% | 35 | 76.09% | 0.59 |
|  | T4 | 23 | 27.38% | 12 | 31.58% | 11 | 23.91% |  |
| Histological grade | High | 10 | 11.90% | 2 | 5.26% | 8 | 17.39% | 0.10 |
|  | Low | 74 | 88.10% | 36 | 94.74% | 38 | 82.61% |  |
| Microsatellite status | MSS | 74 | 88.10% | 36 | 94.74% | 38 | 82.61% | 0.10 |
|  | MSI | 10 | 11.90% | 2 | 5.26% | 8 | 17.39% |  |
| Lymphovascular or perineural invasion | Yes | 13 | 15.48% | 10 | 26.32% | 3 | 6.52% | **0.016** |
|  | No | 71 | 84.52% | 28 | 73.68% | 43 | 93.48% |  |
| Infiltrating margin | Invasive | 49 | 58.33% | 30 | 78.95% | 19 | 41.30% | **0.0008** |
|  | Pushing | 35 | 41.67% | 8 | 21.05% | 27 | 58.70% |  |
| Tumor budding score | Bd1 | 42 | 50% | 13 | 34.21% | 29 | 63.04% | **0.019** |
|  | Bd2 | 16 | 19.05% | 8 | 21.05% | 8 | 17.39% |  |
|  | Bd3 | 26 | 30.95% | 17 | 44.74% | 9 | 19.57% |  |
| Poorly differentiated clusters | G1 | 47 | 55.95% | 15 | 39.47% | 32 | 69.56% | **0.008** |
|  | G2 | 24 | 28.57% | 17 | 44.74% | 7 | 15.22% |  |
|  | G3 | 13 | 15.48% | 6 | 15.79% | 7 | 15.22% |  |
| Lymphocytic CD8+ infiltration | High | 50 | 59.52% | 14 | 36.84% | 36 | 78.26% | **0.0003** |
|  | Low | 34 | 40.48% | 24 | 63.16% | 10 | 21.74% |  |
| PDL1 expression score ($\geq 5\%$) | High | 21 | 25% | 7 | 18.42% | 14 | 30.43% | 0.31 |
|  | Low | 63 | 75% | 31 | 81.58% | 32 | 69.57% |  |
| CDX2 expression | Positive | 81 | 96.43% | 36 | 94.74% | 45 | 97.83% | 0.59 |
|  | Negative | 3 | 3.57% | 2 | 5.26% | 1 | 2.17% |  |

MSS, microsatellites-stable
MSI, microsatellites-instable
PD-L1, programmed death-ligand 1
*P* values $\leq 0.05$ are displayed in bold.

To investigate the independent prognostic value of clinicopathological features, Cox proportional hazards models were fitted for TTR (Table 2). The mean time upon relapse in our cohort was of 1.7 years (range, 0.26 to 5.9 years). The strongest independent predictors in the multivariable setting were CD8+ lymphocytic infiltration (*P* = 0.00019; hazard ratio (HR), 0.25; 95% CI, 0.12 – 0.52) (Supplementary Figures S1B) and lymphovascular or perineural invasion (*P* = 0.0007; HR, 3.70; 95% CI, 1.73 – 7.91) (Supplementary Figures S1C). Additionally, increased tumor budding was also associated with shorter TTR (*P* = 0.026; HR, 1.56; 95% CI, 1.05 - 2.31) (Supplementary Figure S1D).

**Table 2.**
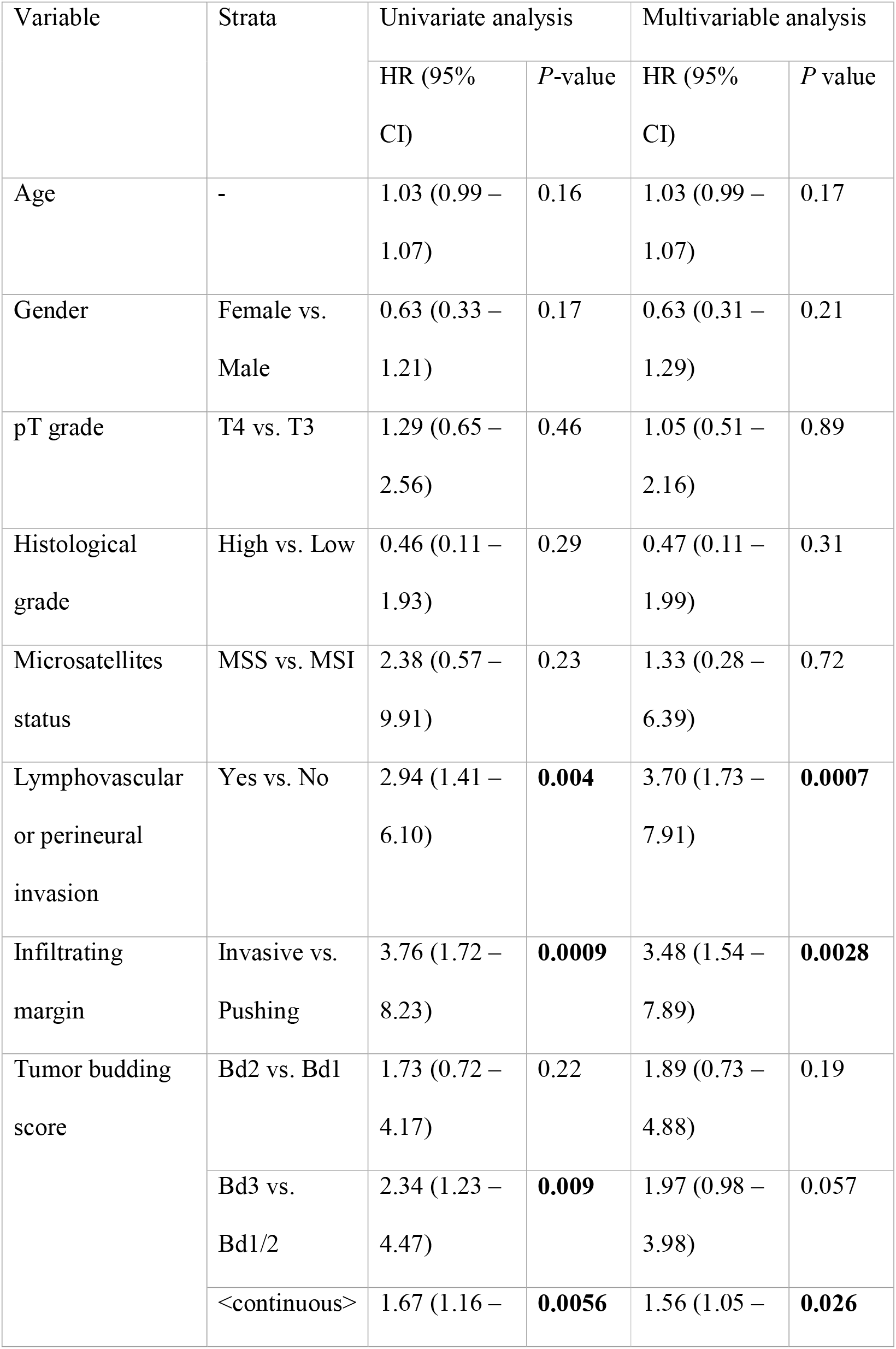
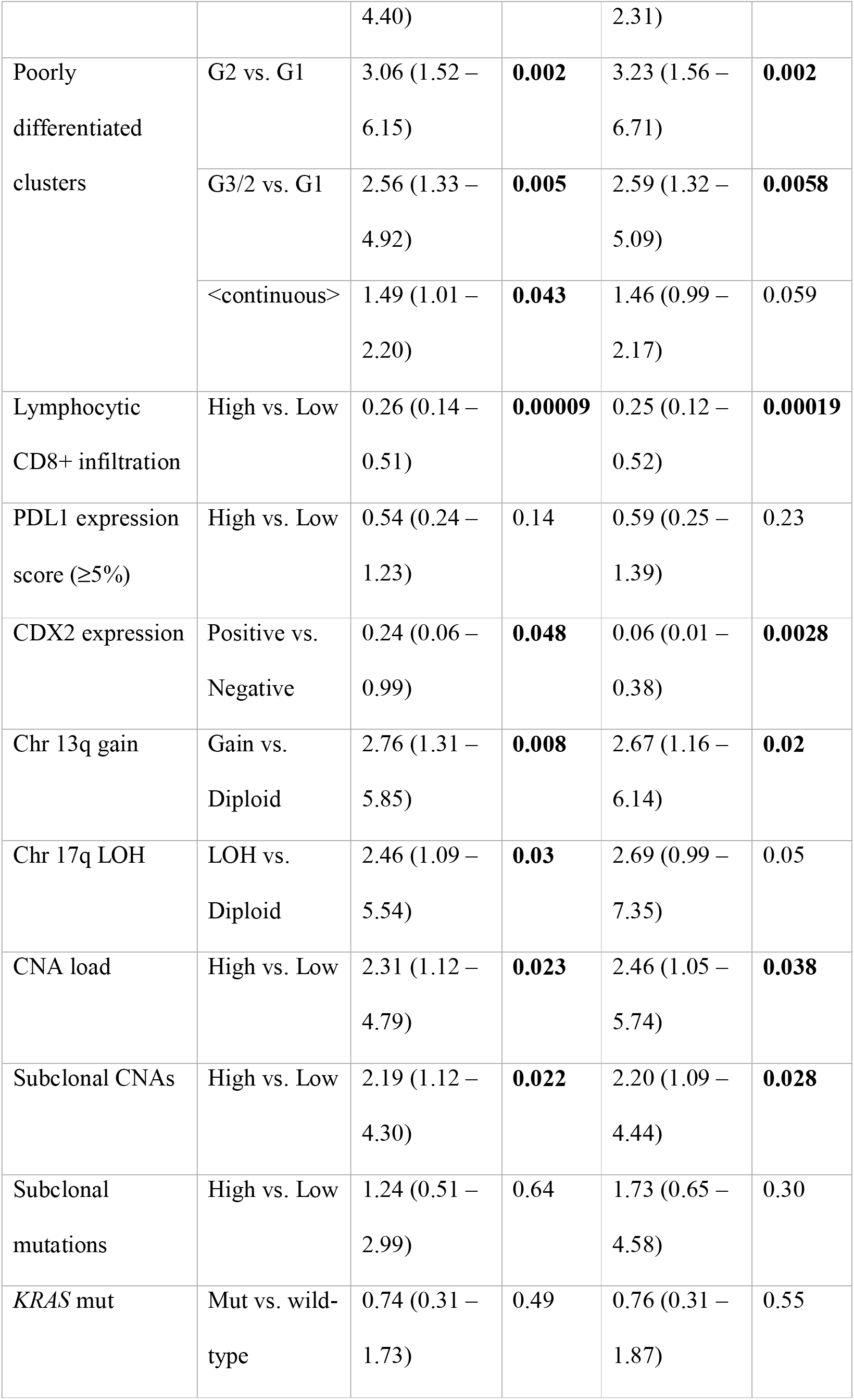
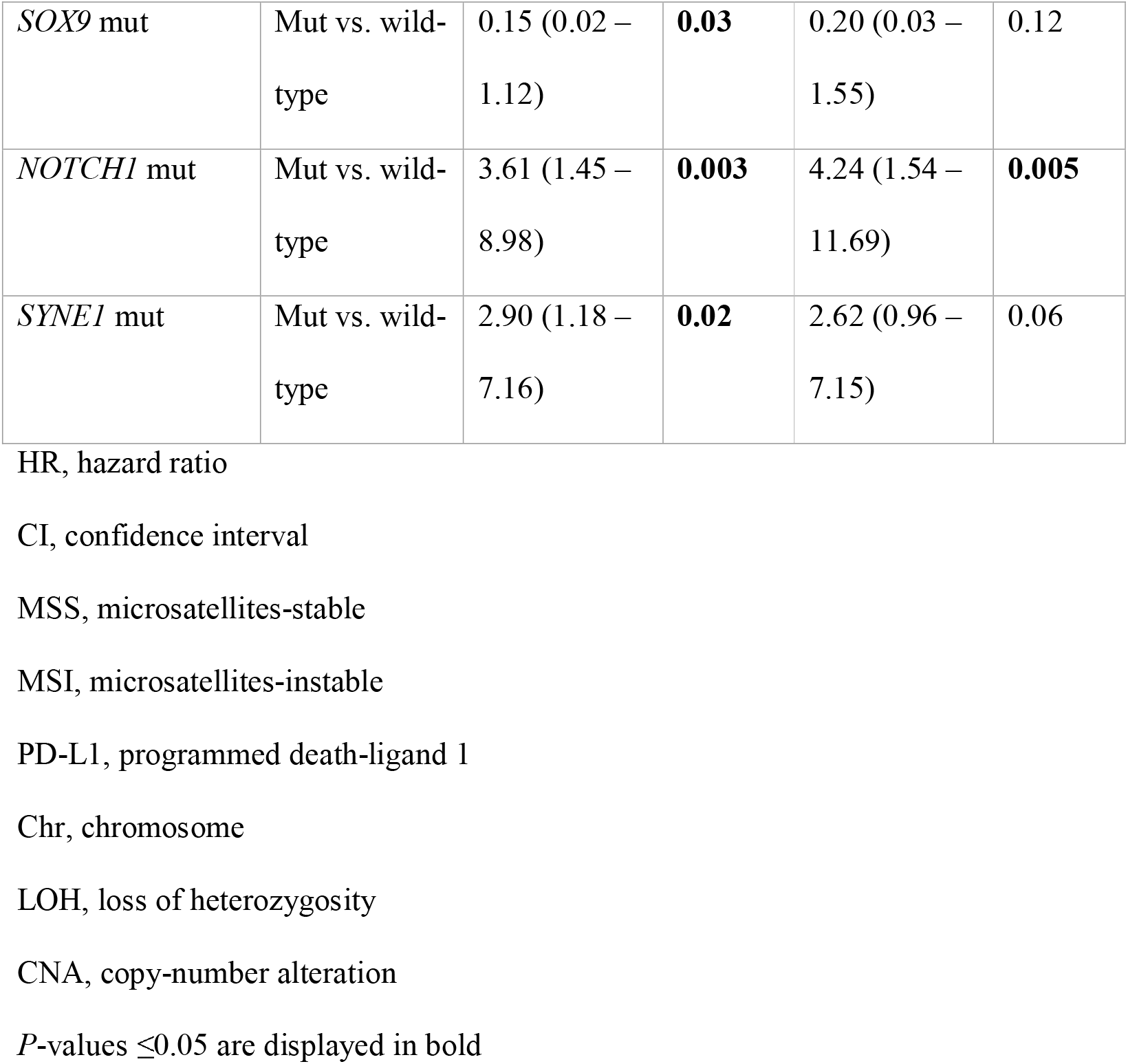
Univariate and multivariable Cox models for TTR, stratified by annotated clinicopathological and tumor-genomic variables.

### Genome-wide screening reveals specific chromosomal aberrations and high CNA load in recurrent tumors

Genome-wide profiling of CNAs was performed in order to identify potential chromosomal regions differentially altered in recurrent versus non-recurrent tumors. Most frequent (>35%) CNAs across all tumors included gains on chromosomes 7, 8q, 13 and 20, and losses (>20%) affecting chromosome arms 5q, 8p, 14q, 15q, 17p and 18q (Figure 1A). The most commonly altered minimal region (<2 Mb) of gain was located at 20q13.32–20q13.33 (71%), and the most repeatedly minimal peak of loss was 18q21.2-18q21.31 (61%).

**Figure 1.**
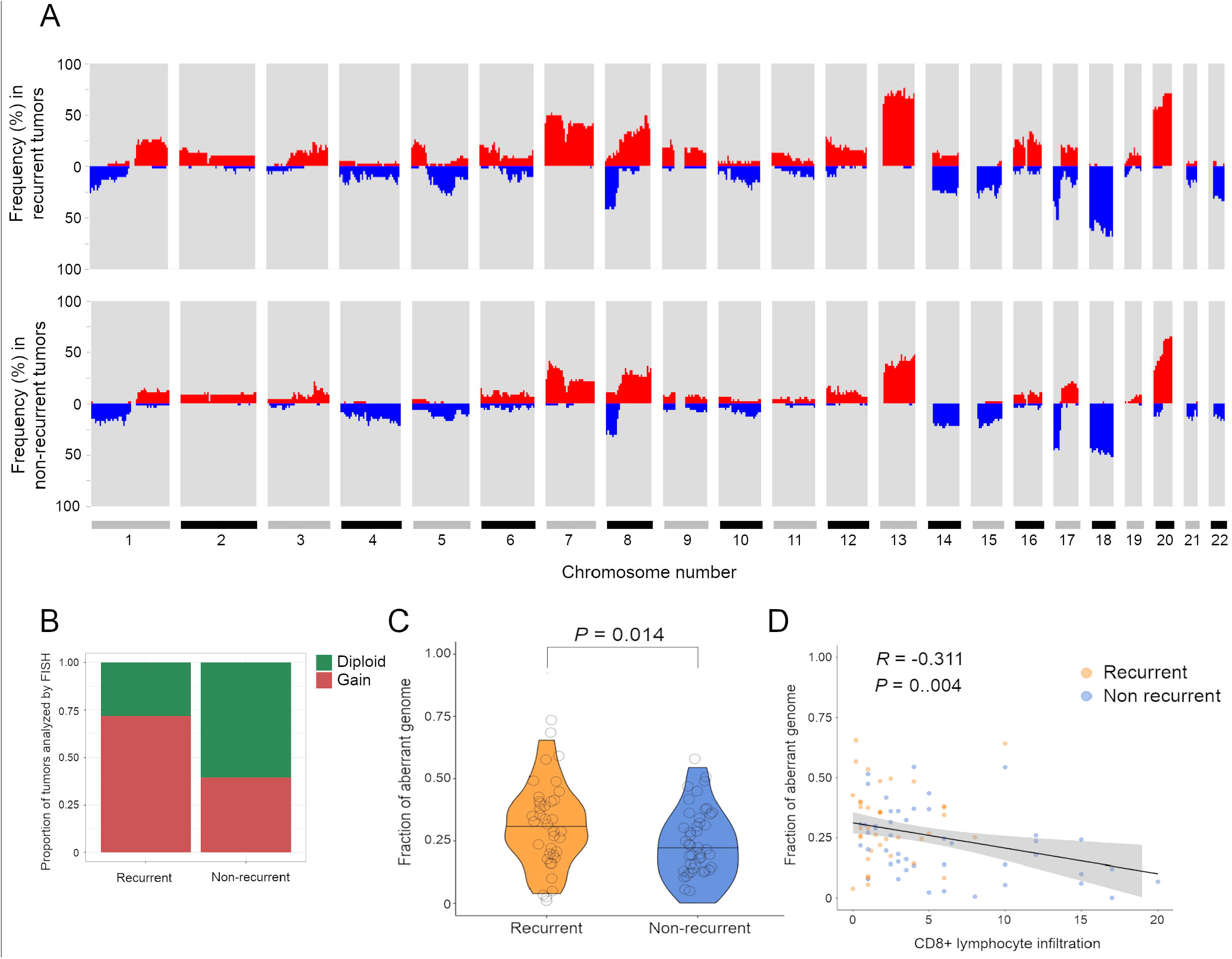
Genome-wide CNA profiles associated with the patient relapse outcome and the tumor CD8+ immune infiltration levels. (A) Whole-genome plot showing the accumulated frequency of copy-number alteration per chromosome, in recurrent (N = 38) and non-recurrent (N = 46) patients. (B) Proportion of tumors with gain (red) or diploid (green) status of the chromosome arm 13q detected by FISH using a surrogate probe covering *CDX2*. (C) Violin plot displaying measures of the fraction of aberrant genome, as a proxy of each tumor level of aneuploidy, in the two clinical groups. Black lines indicate median values. *P* value was obtained by a two-tailed Student’s *t* test. (D) Correlation paired analysis between the proportion of aberrant genome and levels of intratumor CD8+ lymphocytes analyzed by immunohistochemistry. *Rho* and *P* value were obtained with the Spearman’s association test.

Several chromosomal regions appeared significantly more altered in tumors from recurrent patients compared with non-recurrent (Supplementary Table S2). Chromosome arm 13q11-q34 showed the most differential frequency of alteration between the two clinical groups (65% vs. 36% of gains; *P* = 0.018). Two focal CNAs were exclusively present in recurrent tumors, namely the loss of 17q22-q24.3 (12% vs. 0%; *P* = 0.029) and 14q gain (14% vs. 0%; *P* = 0.017) (Figure 1A). In addition, CNN-LOH on 17p13.3-p13.1 was also over-represented in recurrent tumors (23% vs. 5%; *P* = 0.02). Simple linear regression analyses showed a significantly positive correlation between median FISH signals and log2-ratios from SNP-arrays for the two experimentally analyzed aberrations (*R* = 0.82 and *R* = 0.80 for chromosome arms 13q and 17q, respectively; *P* < 0.0001 in both cases) (Supplementary Figure S2A and S2B), thus reproducing the frequencies of CNAs detected by SNP-arrays (Figure 1B).

Since aneuploidy often experiences an increase during cancer progression, the fraction of aberrant genome (hereafter referred as to CNA load) was derived from each tumor SNP-array segmented data. The mean CNA load across tumors was 27%, ranging from 0% to 66%. Tumors from patients with relapse exhibited significantly greater CNA loads in comparison to those without relapse, with mean values being 31.3% in recurrent versus 23% in non-recurrent (*P* = 0.014) (Figure 1C). When excluding MSI positive cases, the mean values were 32.7% versus 26.8%, respectively (*P* = 0.075) (Supplementary Figure S2C). These results were confirmed in the Colonomics cohort, which consisted of 99 MSS stage II colon cancer patients with means of 34.4% in recurrent versus 23% in non-recurrent tumors (*P* = 0.058) (Supplementary Figure S2D). In agreement with previous findings of a decreased cytotoxic immune activity by CD8+ T cells in highly aneuploid tumors [31], we also detected a negative correlation between the CNA load and CD8+ lymphocyte infiltration (*R* = –0.31; *P* = 0.004) (Figure 1D).

### Recurrent tumors exhibit increased levels of subclonal copy-number alterations

Next, we sought to investigate the association of copy-number intratumor heterogeneity with disease relapse. We found considerable levels of subclonality amongst tumors, with 53% of total CNAs appearing as subclonal events, ranging from 0% to 100% per sample. Of those CNAs designated as subclonal, median CCF was 56% [9.8% – 84.9%]. Relapsed tumors carried significantly more subclonal CNAs than non-relapsed, with a median of 10 versus 7 events per group (two-tailed Mann-Whitney *U* test, *P* = 0.02) (Figure 2A and 2B). Statistical significance persisted unchanged when more stringent CCF cut-offs were tested (Supplementary Figure S3A). In addition, subclonal CNAs appeared to positively correlate with CNA load (*R* = 0.52; *P* < 0.0001) (Supplementary Figure S3B), suggesting that intratumor heterogeneity might result from increased aneuploidy.

**Figure 2.**
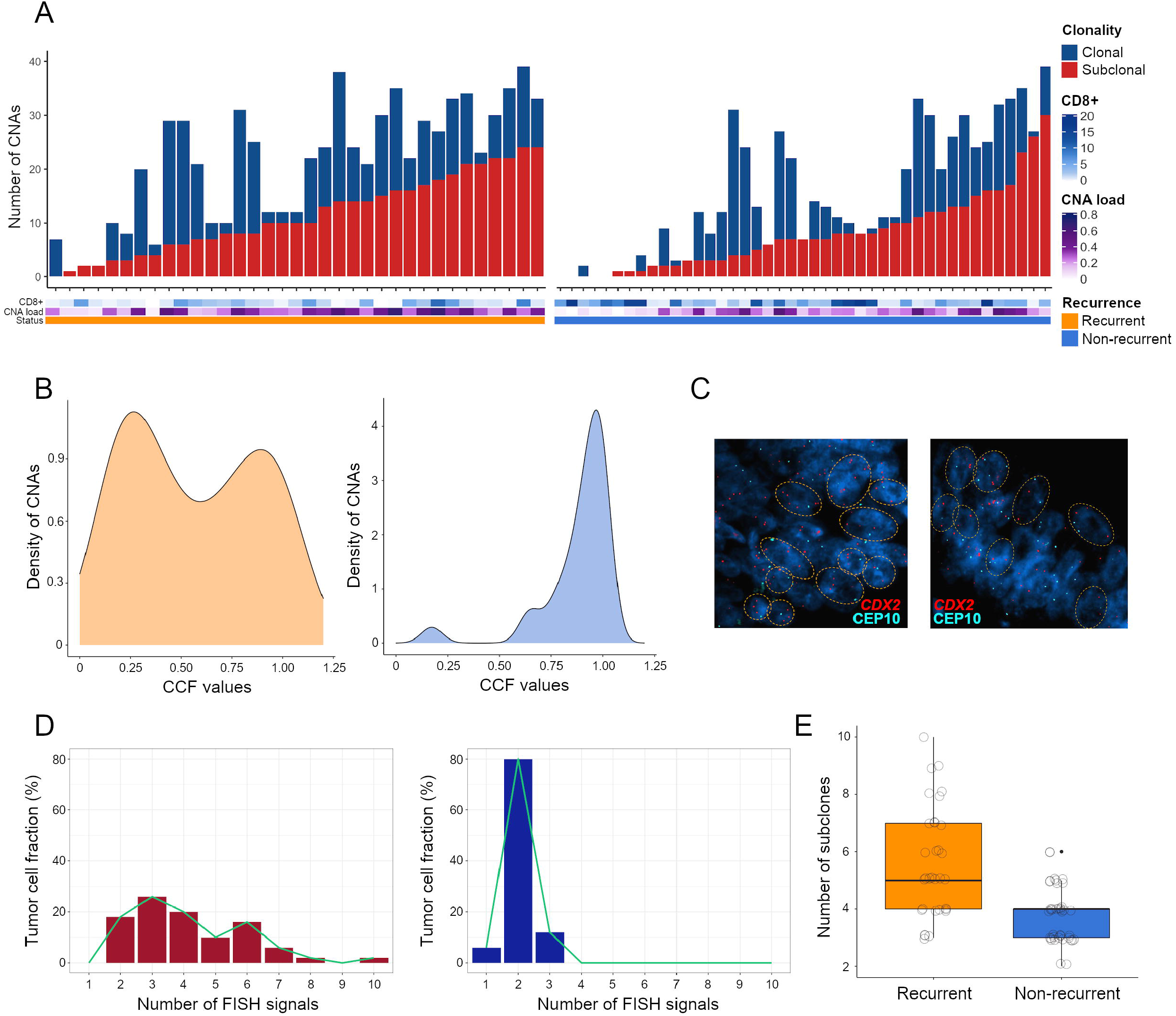
Intratumor heterogeneity of CNAs obtained from untreated patients with stage II colon cancer. (A) Bar plots exhibit the number of CNAs that were found to be clonal or subclonal (<85% of cancer cells) per tumor inferred from SNP-arrays, according to each patient recurrence status, the CNA load and the CD8+ immune infiltration. (B) Representative density plots showing the distribution of CNAs in a recurrent (left panel) and non-recurrent (right panel) tumor. (C) FISH images of a tumor pair with a gained (left) and a diploid (right) 13q, illustrating divergent levels of subclonal heterogeneity. The FISH panel includes fluorescent-labeled probes for *CDX2* (13q) and the centromeric probe CEP10. (D) Histograms depicting the number of cell populations detected by FISH experimental analysis, from two exemplary tumors with different 13q copy-number status and disparate recurrence outcome. Each subclonal population is defined as having a different copy number for the chromosome arm 13q. (E) Box plot exhibits the number of FISH subclonal populations for the chromosome arm 13q, including all tumors from recurrent (N = 38) and non-recurrent (N = 46) patients. *P* value was obtained by a two-tailed Student’s *t* test.

Multi-probe FISH was performed to examine the dynamics of subclonal CNAs for chromosome regions 13q12.2, 17q24.3 and 8q24.21. Results indicated that tumors with copy-number gain on 13q, 17q or 8q exhibited increased subclonal populations compared to those with a diploid status (Supplementary Figure S3C-S3E). Remarkably, when quantifying FISH signals for chromosome arm 13q, relapsed carcinomas displayed increased copy-number diversity in comparison to non-relapsed lesions (*P* < 0.0001) (Figure 2C-2E). In contrast, we did not observe this association for chromosomes 17q nor 8q (Supplementary Figure S3F-S3H), further supporting the selective impact of the chromosome 13q in fostering the subclonal heterogeneity and genomic complexity in early-stage colon tumors.

### Mutational subclonality does not determine disease relapse

To determine whether aneuploidy and levels of subclonal CNAs were associated with the status of the most frequently mutated genes in CRC, a targeted capture DNA sequencing approach was performed on a subset of 44 MSS tumors, including 22 from recurrent patients. Sequencing achieved a mean coverage of 500x in at least 80% of targeted exon regions. Overall, 43 out of the 48 cancer-related genes showed a mutation in at least one patient (Figure 3A). Multi-gene profiling revealed a total of 198 somatic non-synonymous mutations falling in protein-coding regions, accounting for both single-nucleotide variants and small insertions/deletions (Indels) (Supplementary Table S3). Three genes showed a frequency of somatic mutation greater than 20%: *APC* (56.82%), *TP53* (54.55%) and *KRAS* (47.73%) (Figure 3A). As expected, over 85% of mutations affecting *APC* were nonsense or frameshift, while in *TP53* and *KRAS* the majority were missense changes (75% and 100%, respectively). Consistently, 86% of mutations affecting *KRAS* altered hotspot codons 12 or 13, including variant G35A, which represented the 45% of total *RAS* mutations. *SOX9* also manifested a disproportionately high fraction (87.5%) of truncating events.

**Figure 3.**
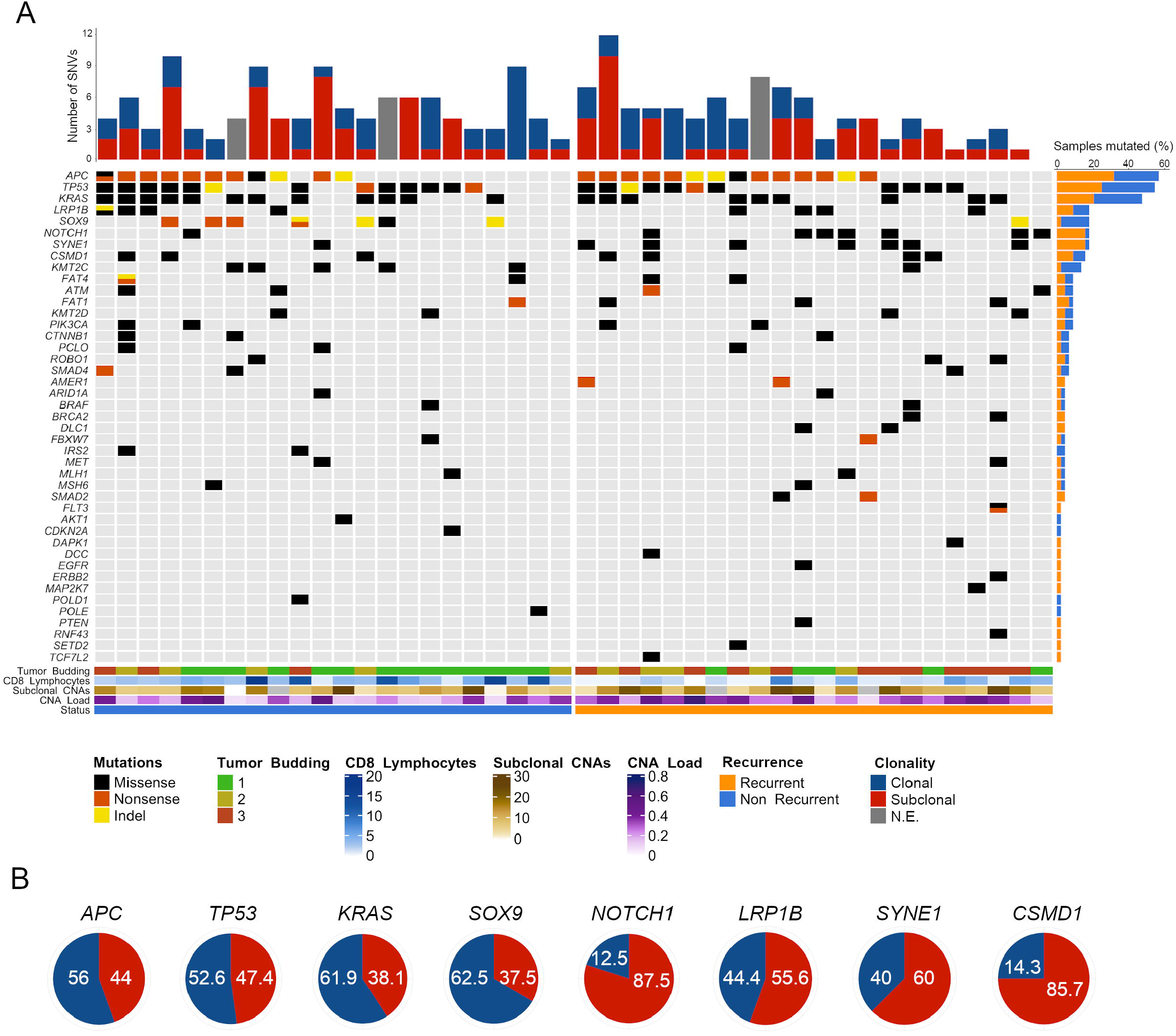
Mutational profiling and subclonal status of MSS tumors from patients with stage II colon cancer. (A) Oncoprint chart illustrates somatic mutation events of the studied genes detected by targeted NGS, arranging cases in two groups according to patient recurrence status. Each column represents an individual tumor and each row a gene. Red/blue bar plot (top) shows the number of mutations per tumor, elucidating their clonal or subclonal (<85%) status. Orange/blue bar plot (right) represents the incidence (number of times) at which each gene was observed mutated. Genes with no mutations are not shown. Color bar plots (bottom) provide data on the genomic and pathological markers annotated in the legend (right). (B) Proportion of mutations that were identified as clonal (blue) or subclonal (red) for the eight most frequently mutated genes. Values are indicated as percentage (%).

No statistically significant differences were observed for *APC, TP53* and *KRAS* mutational status regarding the recurrence condition (Figure 3A) nor the age, gender and any other clinical or genomic features (including the CNA load and subclonal CNAs). In contrast, the mutational frequency of *SOX9, SYNE1* and *NOTCH1* appeared to significantly correlate with recurrence (*P* = 0.046), being *SOX9* more frequently mutated in non-recurrent tumors.

To investigate levels of intratumor heterogeneity concerning single-nucleotide variants, we estimated the CCF harboring each mutation, allowing for discrimination of subclonal (< 85%) from clonal variants. Of the total variants identified, 53% were subclonal, showing a median CCF of 49% [10 to 84%]. In our analysis, recurrent tumors did not exhibit increased subclonal mutations compared to non-recurrent (*P* = 0.75) (Figure 3A). Among genes with a mutational frequency above 15%, *APC, TP53, KRAS* as well as *SOX9* appeared to be mutated in a clonal state in most cases (>50%). Conversely, *NOTCH1, LRP1B, SYNE1* and *CSMD1* were predominantly subclonal (Figure 3B).

### Tumor aneuploidy and intratumor heterogeneity are associated with risk of disease relapse

When assessing the independent prognostic value of the above-described genomic features for TTR (Table 2) (Supplementary Figure S4A), patients with gain of chromosome arm 13q exhibited shorter TTR and a higher risk for recurrence compared to those with a diploid 13q (*P* = 0.02; HR, 2.67; 95% CI, 1.16 – 6.14) in a multivariable analysis correcting for age, sex, pT3/T4, histological grade and microsatellite status (Figure 4A). TTR was also consistently shorter in those patients with tumors carrying LOH at 17q22-q24.3 (i.e., loss or CNN-LOH) (*P* = 0.05; HR, 2.69; 95% CI, 0.99 – 7.35) (Figure 4B).

**Figure 4.**
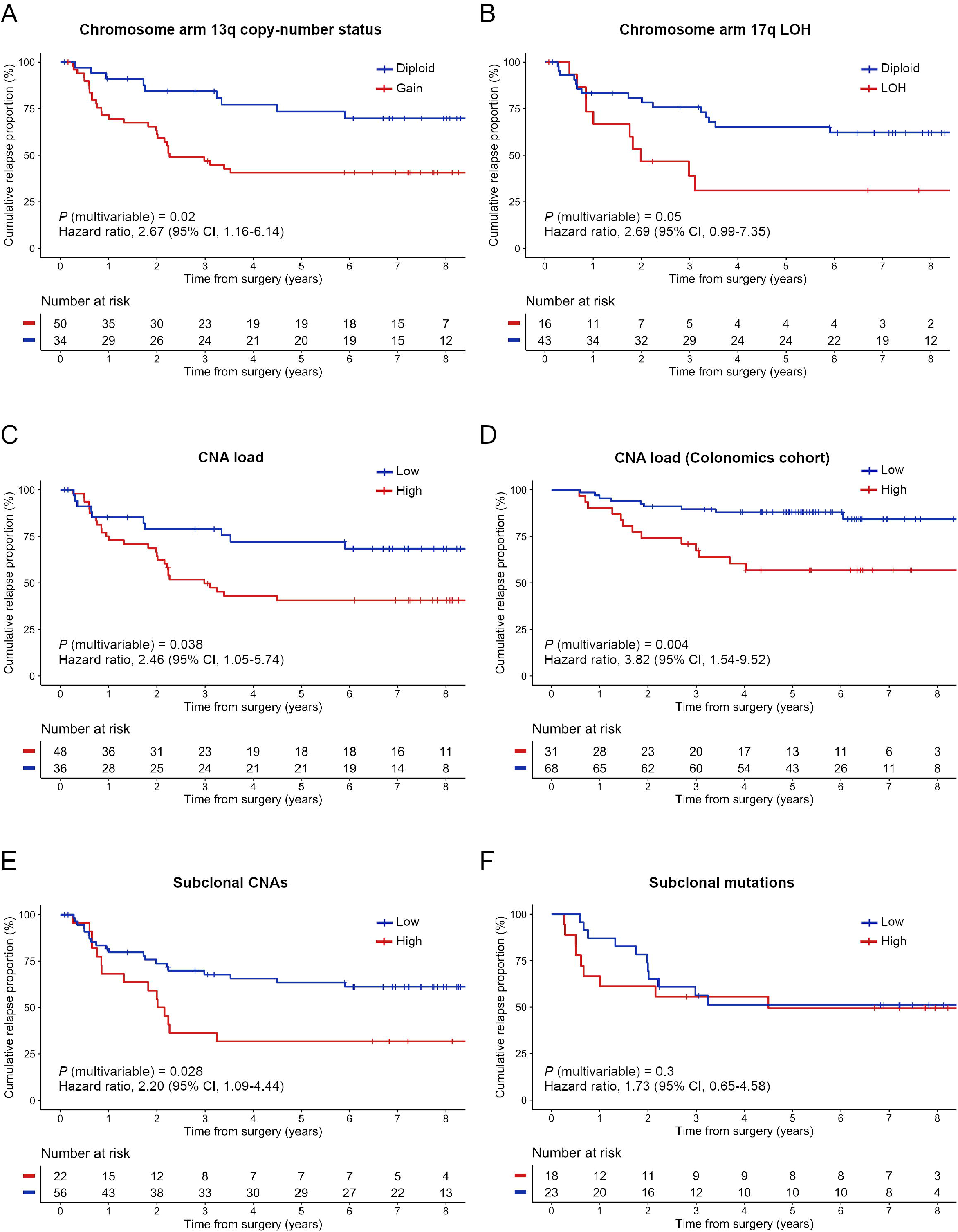
Prognostic value of tumor aneuploidy and genomic heterogeneity for TTR in stage II colon cancer patients. Kaplan-Meier estimates displaying the cumulative proportion (Y axis) of study patients who were relapse-free over a 8-year period (X axis), stratifying by (A) the presence or absence of chromosome arm 13q aberration, (B) 17q22-24.3 LOH (comprising loss or somatic CNN-LOH), (C) a high or low proportion of CNA load in the study population (N = 84), and (D) in an independent cohort of patients (N = 99) diagnosed with MSS tumors, (E) the level of subclonal CNAs identified with SNP-arrays, and (F) a high or low level of subclonal mutations detected by targeted NGS. *P* values were obtained using the log-rank test and hazard ratios using a Cox regression model with proportional hazards, correcting by age, sex, stage pT3/T4, histological grade and microsatellite status.

We then addressed the association of high levels of tumor aneuploidy and genomic heterogeneity with adverse clinical outcome. Kaplan-Meier analysis showed that patients with tumors bearing high CNA loads were at a significantly higher risk of relapse compared to those with low aneuploidy, independently of the patient clinical status (*P* = 0.038; HR, 2.46; 95% CI, 1.05 – 5.74) (Figures 4C). Median time until relapse was 31.48 months in the higher risk group versus 75.28 months in the lower risk. This same association was validated using the independent cohort Colonomics (*P*= 0.004; HR, 3.82; 95% CI, 1.54 – 9.52) (Figure 4D).

Furthermore, elevated copy-number heterogeneity was associated with superior risk of recurrence (*P* = 0.028; HR, 2.20; 95% CI, 1.09 – 4.44) (Figure 4E). The median time until relapse was 25.03 months in the high subclonal CNAs group compared with 70.80 months in the lower group. However, there was no significant correlation of the proportion of subclonal mutations with risk of relapse (*P* = 0.30; HR, 1.73; 95% CI, 0.65 – 4.58) (Figure 4F). Regarding the prognostic value of specific mutated genes, patients with *SOX9* mutations exhibited significantly longer TTR in a univariate analysis (*P* = 0.028; HR, 0.15; 95% CI, 0.02 – 1.12) (Supplementary Figure S4B). Conversely, *NOTCH1* and *SYNE1* mutations were associated with an increased risk of disease recurrence (Supplementary Figures S4C and S4D).

### Genomic markers improve the predictive performance compared to clinicopathological parameters alone

Predictive modeling was performed to classify patients with the primary endpoint of predicting their individual risk to relapse. Genomic and mutational features with the highest prognostic values (Supplementary Figure S5), together with all clinicopathological markers, were included for conventional machine learning analysis to assess their integrated discriminatory power in our cohort of 84 patients. Among the eight classifier algorithms tested (see Supplementary Methods), the gradient boosting machine method was selected as it ranked the shortest fraction of misclassified cases (data not shown). We constructed 4-fold cross-validated models using three different combinations of risk predictors: clinical-, genomic- and clinico-genomic-based, each one comprising the annotated variables indicated in Supplementary Figure S6. For the clinical-based model, the classifier algorithm correctly predicted 56/84 cases (67% accuracy), achieving 64% sensitivity and 69% specificity; for the genomic-based, it rightly spotted 52/84 patients (62% accuracy) with 57% sensitivity and 67% specificity; while for the clinico-genomic model, predictions yielded to 65/84 correctly classified patients (77% accuracy), attaining 73% sensitivity and 81% specificity. For instance, false-positive rates were always inferior to their respective false-negative counts across the three biomarker models (Figure 5A).

**Figure 5.**
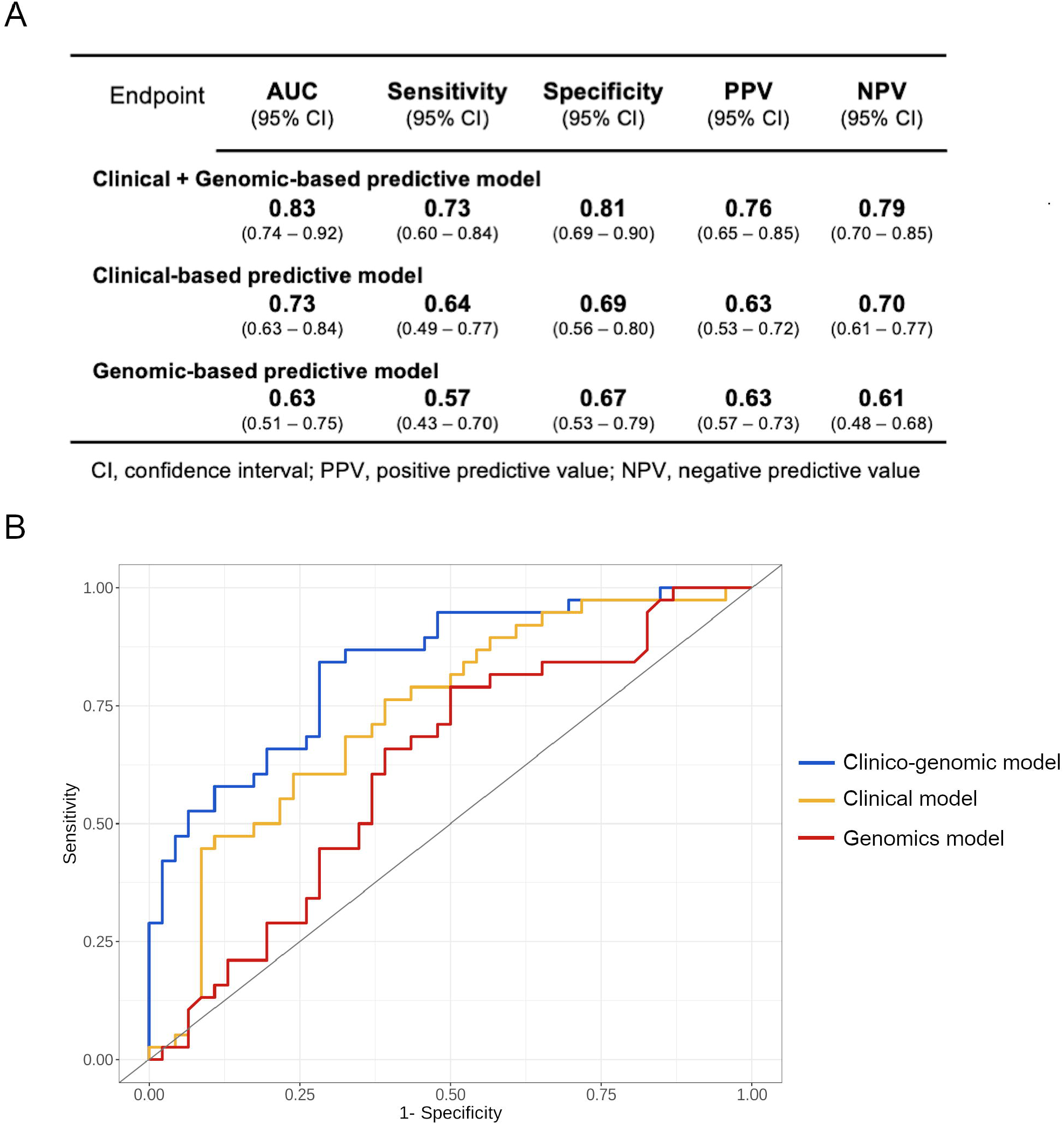
Discriminative ability of the three stage II biomarker models for predicting each patient probability of recurrence. (A) Table showing performance metrics for the three models (i.e., AUC scores, sensibility, specificity, and positive and negative predictive values), obtained by machine learning-based prediction using a gradient boosting machine classifier algorithm. (B) ROC curves are shown for comparative analysis of the three different combinations of variables.

In terms of discriminatory ability, receiving operating characteristic (ROC) curve analyses revealed that the clinico-genomic combination achieved an AUC of 0.83 (95% CI, 0.74 – 0.92), opposing to inferior AUC values of 0.73 (95% CI, 0.63 – 0.84) for the clinical-based and 0.63 (95% CI, 0.51 – 0.75) for the genomic-based models, respectively (Figure 5A-B). The addition of the genomic-related variables upon baseline clinicopathological parameters caused a significant 10% incremental gain in the AUC (DeLong’s *P* = 0.03).

## DISCUSSION

Intratumor heterogeneity fosters the evolution of the genome leading to metastatic progress and therapy resistance [8]. In the present study, we investigate the relative contribution of genomic heterogeneity involving CNAs and mutational events as a prognostic determinant for disease recurrence in untreated stage II colon cancer patients. Additionally, we construct a predictive model integrating the levels of genomic complexity along with clinicopathological features, enabling the identification of those patients at high-risk of recurrence with notable discriminative ability.

Tumor aneuploidy, or the presence of chromosome copy-number imbalances, is a hallmark of human solid malignancies [32] and has been depicted as an unfavorable prognostic factor pan-cancer [33,34]. We define the fraction of aberrant genome, or CNA load, as a static measure of whole-genome levels of chromosome disruption. An increased CNA load has already been evidenced to predict poor outcome in metastatic CRC [25], yet has not been thoroughly explored in early-stage disease as a prognostic biomarker. In our cohort of stage II colon cancer, tumors from recurrent patients encompass significantly higher CNA load compared with non-recurrent. This trend is maintained when excluding MSI+ cases, which in general exhibit near-diploid karyotypes and fall in the non-recurrent group. Importantly, our data support a strong correlation between genome aneuploidy and lower probability of being relapse-free in two independent cohorts, with the consideration that the validation cohort consisted of MSS tumors exclusively. In an attempt to identify chromosome-specific regions with ability to discriminate recurrent lesions, we found several CNAs including the gain of chromosome arm 13q and loss at 17q22-q24.3, subsequently confirmed by FISH. In line with this, a previous study already showed that gains of *CDX2* were exclusively seen in primary recurrent adenomas compared to those without recurrence [35]. Conclusively, these findings might provide evidence on the prognostic contribution of tumor aneuploidy in early-stage colon cancer.

To capture an approximative measure of the degree of intratumor heterogeneity of each tumor, we determined the subclonal status of CNAs, CNN-LOH and mutations. The fact that all patients included in this study were chemotherapy-untreated avoids the genetic bottleneck caused by cancer therapy enabling a better preservation of the original overall clonal diversity. Increased levels of copy-number subclonality have been associated with poor outcome in non-small cell lung cancer [36] and in hepatocellular carcinoma [20], while mutational multi-clonality correlated with worse disease-free survival in stages I-IV CRC [37]. Likewise, our data show that tumors with elevated subclonal copy-number dosage exhibit significantly shorter TTR, making plausible that early-stage tumors with extensive copy-number heterogeneity might fall in the high-risk group susceptible to benefit from adjuvant chemotherapy. Besides, we find that tumors with high CNA load display increased degrees of copy-number subclonality, which reinforces the idea that intratumor heterogeneity predominantly occurs in highly aneuploid tumors through a process governed by ongoing CIN. In contrast, the amount of subclonal mutations did not appear to be associated neither with recurrence in our cohort, strengthening the potential prognostic power of CIN over the tumor mutational burden, possibly because a sole copy-number alteration can disrupt the transcription of a multitude of genes simultaneously [38,39]. Our results are consistent with previous reports showing that gene copy numbers, but not coding mutations, are highly discordant between colorectal primary tumors and their matched metastases [40]. In light of these results, the need of assessing CNAs using noninvasive approaches, such as liquid biopsy, urges further attention in the clinical setting.

The mutational landscape of CRC has been well-characterized [41]. Our results suggest that copy-number subclonality is independent from the mutational status of the main CRC driver genes (i.e., *APC, TP53* and *KRAS*). We find that none of the previous three mutated genes correlated with risk of relapse, in concordance with reported data in stages II-III colon cancer [4,42]. Noteworthy, the mutational status of *SOX9, NOTCH1* and *SYNE1* displayed prognostic value in our cohort. In CRC, overexpression of NOTCH1 has been appointed as a negative predictor of overall survival [43], consistent with our findings for TTR. Conversely, patients with mutations in *SOX9* exhibited longer TTR in our series, but this association appears to weaken in the multivariable setting. This result is in line with a previously reported association of *SOX9* mutations with increased overall survival in metastatic CRC [44] and overexpression of SOX9 at the invasive front with significantly higher relapse-free times in stage II colon cancer [45]. Finally, our observations on *SOX9*-mutated and *APC*-wildtype tumors displaying more abundant CD8+ lymphocyte populations than their counterparts suggest a putative role of the WNT/β-catenin pathway in modulating the tumor-infiltrating T cell compartment [46].

Besides the independent prognostic value of genomic markers, our data reinforce the idea that tumor microenvironment features (e.g., low intratumor CD8+ lymphocytic infiltration) and the presence of lymphovascular or perineural invasion are still the most potent determinants of poor outcome in locally advanced colon cancer. The impact of anticancer immune cytotoxicity as a major determinant for disease recurrence in stages II-III has been widely validated in multiple independent series of patients over the past 5 years [4,47,48]. In addition, tumor budding and poorly differentiated clusters might constitute early histological features potentially leading to an ulterior metastatic phenotype [5], which might explain their strong ability to predict risk of relapse in early-stage colon cancer. Finally, in a large-cohort study, the lack of CDX2 expression was associated with shorter relapse-free survival in stage II colon cancer [49]. We observed the same trend regardless the low number of tumors with CDX2 negative expression.

Despite the plethora of identified prognostic biomarkers, prediction of stage II colon cancer recurrence based on molecular information is still an open problem. Here, we created a bioinformatic machine learning approach integrating a range of twenty-two risk predictors. The best configuration of our clinico-genomic model achieved significant discriminant capacity, rating an AUC of 83%, comparable with previous reported AUC values in various stage II colon cancer cohorts using molecular-based signatures [50,51]. In our predictive model, the addition of genomic-related markers to the patient clinicopathological information caused a significant increase in the AUC of 10 points, which was also associated with a similar increase in the sensitivity and accuracy. Intriguingly, across the three combination models, specificity values were always higher than their respective sensitivities by 5-10 points, stressing the need of identifying more sensitive strategies capable of reducing false-negative rates. An obvious limitation is a number of twenty-two predictors included in the clinico-genomic model appears to be unmanageable in any clinical routine.

To summarize, our results reinforce the potential value of intratumor heterogeneity driven by chromosomal instability as a prognostic factor in early-stage colon cancer. In this scenario, patients with tumors harboring high levels of aneuploidy and copy-number subclonality might be at risk of relapse and could benefit from early therapeutic intervention during disease monitoring. Given the retrospective design of this study, we advocate for these results to be further validated in randomized, prospective clinical trials incorporating NGS strategies, intending to optimize patient stratification at the adjuvant setting.

## ACKNOWLEDGEMENTS

The authors would like to thank Dr. Joaquim Radua (from IDIBAPS) for statistical assistance and Dr. Daniel Aguilar (from CIBEREHD) for bioinformatics support. The authors are also thankful to the Biobank-Tumor Bank and Functional Genomics platforms from IDIBAPS-Hospital Clínic of Barcelona for preparing and processing samples and for microarrays and next-generation sequencing procedures. The study has been developed in part in the Centre Esther Koplowitz from IDIBAPS / CERCA Programme / Generalitat de Catalunya.

