## Supplemental for "Copy-number intratumor heterogeneity contributes to predict relapse in chemotherapy-naive stage II colon cancer"

**CONTENTS**

**Supplementary Materials and Methods:**

- Cohort design and tissue microarray construction
- SNP-arrays
- Targeted next-generation sequencing (NGS)
- Fluorescence in situ hybridization
- Immunohistochemistry
- Prognostic modeling
- Statistical analysis

**References**

**Supplementary Figure Legends**

**Supplementary Tables**

**Supplementary Figures**

#### SUPPLEMENTARY MATERIALS AND METHODS

##### Cohort design and tissue microarray construction

Eighty-four stage II colon adenocarcinomas (pT3-4N0M0) provided by the Hospital Clínic of Barcelona/IDIBAPS Tumor Biobank were retrospectively collected for this study. The study cohort was designed to maintain a balanced ~1:1 distribution of cases with and without relapse, aimed at maximizing the statistical power for ulterior statistical analysis. Samples were obtained from formalin-fixed paraffin-embedded (FFPE) tissue blocks after macrodissection by an expert gastrointestinal pathologist to minimize inclusion of non-desired tissue components such as necrotic tissue and normal mucosa. Tissue microarrays (TMA) were constructed including tissue sections of 1 mm-core diameter and 4 µm of thickness. Each TMA comprised one replicate of normal adjacent mucosa, four replicates of the bulk primary tumor, and four replicates of the tumor infiltrating margin for each case. Sections of each TMA were used for FISH and immunohistochemistry stains.

##### SNP-arrays

DNA was extracted from each sample using the QIAmp DNA FFPE kit (Qiagen, Hilden, Germany) and then quantified using Qubit fluorometer assay (Thermo Fisher Scientific, MA, USA). Two hundred nanograms of DNA were processed for hybridization on arrays following the commercial protocol OncoScan FFPE assay kit based on Molecular Inversion Probe technology (Affymetrix, Santa Clara, CA, USA). After hybridization, microarrays were processed and scanned using GeneChip instrumentation system 3000 from Affymetrix. Samples from the COLONOMICS dataset were previously processed using Affymetrix Genome-Wide Human SNP Array 6.0 genotyping arrays

(EGAD00010001253).

#### Targeted next-generation sequencing (NGS)

Enriched libraries were prepared from 50 ng of DNA per tumor sample using the SMARTer ThruPLEX DNA-seq Library Preparation kit (TaKaRa, Japan) and a predesigned gene capture pool of IDT xGen Lockdown probes (IDT, Coralville, Iowa, USA). Paired-end sequencing runs were performed on a MiSeq platform (Illumina, San Diego, CA, USA) following a protocol with 2x130 bp reads.

#### Fluorescence *in situ* hybridization

The following BAC clones were used to generate FISH probes: one covering *CDX2* on 13q12.2 (RP11-153M24), another located on *SOX9* on 17q24.3 (RP11-727K24) and a third one covering *MYC* gene on 8q24.21 (CTD-3056O22), all tagged with Spectrum Red-dUTP (Enzo Life Sciences, NY, USA) following a standard nick-translation protocol. Three centromeric FISH probes (CEP) were used as positive controls for chromosomes 6, 10 and 17 labelled with Spectrum-Aqua (Abbot Molecular, Illinois, USA). Tumor sections on TMAs went through deparaffination, pepsin regime, sample pre-treatment and post-treatment as described previously [1]. Probes were denatured at 85ºC during 3-5 minutes and hybridized at 37ºC overnight in a Dako Hybridizer (Agilent Technologies, Santa Clara, CA, USA). Between 50-150 interphasic nuclei per tumor were imaged with a Nikon Eclipse 50i fluorescence microscope using the Isis Fluorescence Imaging System (MetaSystems, Altlussheim, Germany). All countings were done in a blinded-fashion manner by three independent researchers (SL, EA and JdR). A cell population with a discrete number of FISH signals was considered to be a subclone if it was present in at least 2% of aberrant cells detected by FISH.

**Immunohistochemistry**

Immunohistochemical stains were performed on 4 μm TMA cores or on whole tumor sections, using the Ventana Benchmark automated staining system (Ventana Medical Systems, Tucson, USA) based on the manufacturer's recommendations. Antibodies used for stainings were: CDX-2 (EPR2764Y, 760-4380, Cell Marque), CD8 (SP57, 790-4460, Roche), mismatch repair (MMR) proteins including MLH1 (ES05, IR079, Dako), PMS2 (EP51, IR087, Dako), MSH2 (G219-1129, 790-5093, Cell Marque) and MSH6 (SP93, 790-5092, Roche), PD-L1 (22C3, pharmDx), and CKAE1/AE3 staining (AE1+AE3, PDM072, Diagnostic BioSystems). As positive controls, we used placenta for PD-L1, tonsil for CD8, and normal appendix for CDX-2, CKAE1/AE3, MLH1, MSH2, PMS2, and MSH6. Negative controls were carried out by omitting the primary antibody. CD8 immune infiltration was considered high when there were more than 2 tumor infiltrating lymphocytes (TIL) per high-power field x400 (HPF) after counting 5 non-contiguous 40x fields and calculating the mean number of TIL/HPF. Tumors were classified as microsatellite-stable (MSS) when tumor cells presented nuclear staining on the four MMR antibodies; instead, they were considered as microsatellite-instable (MSI) when the tumor cell nuclei were negative for one or more antibodies. Nuclear staining of stromal cells and lymphocytes served as an internal positive control. PD-L1 was scored in whole tissue sections using the combined positive score (CPS), defined as the number of positive tumor cells, lymphocytes and macrophages, divided by the total number of viable tumor cells multiplied by 100, and was considered positive when >5% positive cells. Tumor budding and poorly differentiated clusters were scored on whole tissue sections using CKAE1-AE3 staining, and were graded on a three-tier system [2,3]. Immunostainings were assessed blindly by two independent pathologists (IA and MC) using a BX51 microscope (Olympus, Tokyo, Japan). Discrepancies were resolved by simultaneous re-evaluation, and a consensus decision was made.

**Prognostic modeling**

The Kaplan-Meier method from *survival* R package was used to estimate time-point curves, and the log-rank was applied to assess statistical differences on time to recurrence (TTR) between arms. Hazard ratios and associated 95% confidence intervals were calculated by univariate and multivariable Cox regression models with proportional hazards. Assumption of hazards linearity was tested by checking the Schoenfeld residuals of Cox models ($P < 0.05$). Multivariable models were adjusted by age, sex, pT3/T4, histological grade and microsatellite status.

**Statistical analysis**

All calculations were performed with R statistical software version 3.6. Normality of continuous data was tested using Shapiro-Wilk's statistics. To check for associations between groups, Student's *t*-test and Mann-Whitney-Wilcoxon rank-sum test were used for parametric and non-parametric data, respectively. Fisher's exact or Chi-squared tests were employed in case of categorical classes. To assess statistical correlation of continuous variables, Pearson or Spearman's tests were applied for parametric and non-parametric, respectively. Cut-offs to categorize continuous variables were calculated using functions from R-packages *survMisc* and *pROC*, unless indicated another method. All $P$ values were double-sided and considered significant when < 0.05.

#### SUPPLEMENTARY FIGURE LEGENDS

**Figure S1: Kaplan-Meier estimates on TTR stratified by the annotated immunohistochemistry-based variables.** (A) Representative images of a high (top) and low (bottom) CD8+ lymphocyte infiltration tumor. Scale bar, 100 µm. Relationship of TTR with (B) levels of CD8+ lymphocytic infiltration, (C) a high or low proportion of lymphovascular or perineural invasion by the tumor cells, and (D) tumor budding categories. Cox regression models were fitted to obtain multivariable hazard ratios, adjusting by age, sex, stage pT3/T4, histological grade and microsatellite status, and the log-rank test to obtain *P* values.

**Figure S2: Correlation of FISH counts with SNP-arrays copy-number quantification.** Scatter plots and simple linear regression analysis of log2 ratio values (from SNP-arrays) with the median number of FISH signals for (A) the chromosome arm 13q, using a probe for *CDX2* (N=82), and for (B) the chromosome arm 17q, using a probe for *SOX9* (in N=76). Violin plots showing measurements of the CNA load in tumors from recurrent and non-recurrent patients, excluding cases with microsatellite-instability (MSI) in (C) our own cohort and in (D) the Colonomics cohort. Black lines indicate median values, and *P* values were obtained by a Student's *t* test.

**Figure S3: Extended assessments to validate intratumor heterogeneity analysis.** (A) Graphical representation of the variation of the *P* value in relation to the different cut-offs tested for the cancer cell fraction (CCF) to consider a CNA event as clonal or subclonal when comparing between recurrent and non-recurrent tumors. (B) Density plot and simple linear regression analysis of the number of subclonal CNAs (Y-axis) relative to the CNA load (X-axis). Colors indicate the distribution of tumors according to their recurrent/non-recurrent status. *Rho* and *P* value were obtained with a Spearman's association test. Box plots exhibiting numbers of FISH subclonal populations for chromosome arms (C) 13q, (D) 17q and (E) 8q, in tumor groups with and without CNA affecting these chromosomes. *P* values were derived from a two-tailed Student's *t* test. Box plot exhibiting the number of FISH subclonal populations for the chromosome arms (F) 13q, (G) 17q and (H) 8q, grouping tumors whether they are recurrent or non-recurrent.

**Figure S4: Kaplan-Meier measures of TTR stratified by the mutational status of three genes tested by NGS.** (A) Tree plot exhibiting the hazard ratio (HR) for relapse associated with each chromosome specific aberration. Relapse-free probability depending on the mutational status of (B) *SOX9*, (C) *NOTCH1* and (D) *SYNE1*, on the subset of *N*=44 MSS tumors included in the NGS analysis. Cox regression models were fitted to obtain HRs, and the log-rank test to obtain *P* values. Multivariable estimates were corrected by age, sex, stage pT3/T4 and histological grade. Uv., univariate; mv., multivariable.

**Figure S5. Analysis of the prognostic risk of the clinico-genomic variables included in the predictive model.** Tree plot exhibiting HRs concerning all the clinico-genomic variables included in the predictive model. HRs and associated 95% confidence intervals (CI) were inferred from univariate Cox regression models. *P* values were obtained by log-rank statistics.

**Figure S6. Flowchart of the machine learning-based approach to develop our predictive model based on the annotated clinical and genomic variables.**

**Table S1. Summary of the targeted genes and exon regions tested in the NGS analysis**

| Gene | Chrom | Targeted_Regions | Size_bp | Percent_Covered | Seq_Start | Seq_Stop | Strand |
|---|---|---|---|---|---|---|---|
| AKT1 | 14 | 14 | 1615 | 100 | 105236678 | 105258980 | - |
| AMER1 | X | 3 | 7170 | 97.98 | 63405997 | 63413166 | - |
| APC | 5 | 19 | 8931 | 100 | 112043415 | 112179823 | + |
| ARID1A | 1 | 24 | 6858 | 99.83 | 27022895 | 27107247 | + |
| ARID2 | 12 | 22 | 5508 | 99.46 | 46123620 | 46298861 | + |
| ATM | 11 | 62 | 9188 | 99.81 | 108098410 | 108236235 | + |
| BRAF | 7 | 20 | 10204 | 98.71 | 140419134 | 140624305 | - |
| BRCA2 | 13 | 26 | 10257 | 100 | 32890598 | 32972907 | + |
| CDKN2A | 9 | 8 | 2627 | 98.48 | 21968228 | 21994357 | - |
| CSMD1 | 8 | 70 | 10695 | 100 | 2796107 | 4851938 | - |
| CTNNB1 | 3 | 14 | 2346 | 100 | 41265560 | 41280833 | + |
| DAPK1 | 9 | 26 | 4400 | 99.95 | 90113993 | 90322279 | + |
| DCC | 18 | 29 | 4344 | 100 | 49867216 | 51057023 | + |
| DLC1 | 8 | 22 | 5992 | 99.68 | 12943320 | 13357580 | - |
| EGFR | 7 | 33 | 4952 | 98.81 | 55086971 | 55273310 | + |
| ERBB2 | 17 | 28 | 4036 | 100 | 37855813 | 37884297 | + |
| FAT1 | 4 | 27 | 13821 | 100 | 187509746 | 187630999 | - |
| FAT4 | 4 | 17 | 14952 | 100 | 126237567 | 126412923 | + |
| FBXW7 | 4 | 16 | 3069 | 98.11 | 153244241 | 153333024 | - |
| FLT3 | 13 | 24 | 2982 | 100 | 28578189 | 28674647 | - |
| GNAS | 20 | 22 | 5539 | 96.71 | 57415162 | 57485884 | + |
| HRAS | 11 | 5 | 699 | 100 | 532589 | 534322 | - |
| IRS2 | 13 | 6 | 4017 | 96.12 | 110408651 | 110438400 | - |
| KMT2C | 7 | 64 | 14958 | 99.74 | 151833917 | 152132871 | - |
| KMT2D | 12 | 59 | 16614 | 99.39 | 49415563 | 49449107 | - |
| KRAS | 12 | 5 | 1273 | 100 | 25362445 | 25398329 | - |
| LRP1B | 2 | 90 | 13800 | 99.28 | 140990755 | 142888239 | - |
| MAP2K7 | 19 | 12 | 1329 | 100 | 7968938 | 7977316 | + |
| MET | 7 | 21 | 4425 | 97.36 | 116339125 | 116436178 | + |
| MLH1 | 3 | 20 | 2538 | 94.29 | 37035039 | 37092144 | + |
| MSH2 | 2 | 17 | 3216 | 100 | 47630331 | 47739573 | + |
| MSH6 | 2 | 12 | 4292 | 100 | 48010373 | 48033999 | + |
| NOTCH1 | 9 | 35 | 7668 | 99.53 | 139390523 | 139440238 | - |
| NRAS | 1 | 4 | 570 | 100 | 115251156 | 115258781 | - |
| PCLO | 7 | 27 | 17444 | 99.72 | 82387891 | 82791908 | - |
| PIK3CA | 3 | 20 | 3207 | 100 | 178916614 | 178952152 | + |
| POLD1 | 19 | 27 | 3402 | 99.62 | 50902109 | 50921204 | + |
| POLE | 12 | 49 | 6861 | 100 | 133201283 | 133263901 | - |
| PTEN | 10 | 9 | 1212 | 100 | 89624226 | 89725229 | + |
| RNF43 | 17 | 9 | 2352 | 100 | 56432304 | 56492938 | - |
| ROBO1 | 3 | 33 | 6018 | 100 | 78648063 | 79639061 | - |
| SETD2 | 3 | 24 | 7978 | 99.32 | 47058583 | 47205414 | - |
| SMAD2 | 18 | 10 | 1451 | 100 | 45368198 | 45423174 | - |
| SMAD4 | 18 | 13 | 1772 | 97.63 | 48573417 | 48604837 | + |
| SOX9 | 17 | 3 | 1530 | 100 | 70117533 | 70120528 | + |
| SYNE1 | 6 | 146 | 26503 | 100 | 152443571 | 152949466 | - |
| TCF7L2 | 10 | 20 | 2998 | 100 | 114710516 | 114925731 | + |
| TP53 | 17 | 10 | 1689 | 92.54 | 7572927 | 7579912 | - |

**Table S2. Chromosomal regions with a differential frequency of copy-number or LOH alteration between relapsed and non-relapsed tumors identified by SNP arrays**

| Chrom | Start | End | Cytob_Start | Cytob_End | Event | Length | Freq_Rec | Freq_NonRec | Freq_Diff | p.value | Genes |
|---|---|---|---|---|---|---|---|---|---|---|---|
| 5 | 11151130 | 45485815 | p15.2 | p12 | Gain | 34334686 | 23.76 | 4.33 | 19.42 | 0.0209 | 144 |
| 6 | 0 | 24911553 | p25.3 | p22.3 | Gain | 24911554 | 24.62 | 6.10 | 18.52 | 0.0238 | 154 |
| 7 | 3477017 | 9126751 | p22.2 | p21.3 | Gain | 5649735 | 41.23 | 18.38 | 22.85 | 0.0291 | 12 |
| 7 | 46052682 | 54235567 | p12.3 | p12.1 | Gain | 8182886 | 51.19 | 28.06 | 23.12 | 0.0408 | 11 |
| 7 | 66849481 | 80168569 | q11.21 | q21.11 | Gain | 13319089 | 36.98 | 14.18 | 22.80 | 0.0240 | 135 |
| 7 | 85183242 | 111394602 | q21.11 | q31.1 | Gain | 26211361 | 43.55 | 19.57 | 23.98 | 0.0258 | 257 |
| 7 | 124391281 | 133317496 | q31.33 | q32.3 | Gain | 8926216 | 41.43 | 17.95 | 23.47 | 0.0239 | 56 |
| 7 | 148572283 | 159138663 | q36.1 | q36.3 | Gain | 10566381 | 41.16 | 16.96 | 24.19 | 0.0194 | 107 |
| 9 | 0 | 11510803 | p24.3 | p23 | Gain | 11510804 | 24.14 | 6.32 | 17.81 | 0.0280 | 33 |
| 9 | 17769612 | 38737234 | p22.2 | p13.2 | Gain | 20967623 | 26.58 | 8.04 | 18.54 | 0.0311 | 63 |
| 11 | 7406221 | 14996292 | p15.4 | p15.2 | Gain | 7590072 | 17.23 | 2.43 | 14.79 | 0.0353 | 37 |
| 13 | 19084823 | 115169878 | q11 | q34 | Gain | 96085056 | 65.27 | 36.41 | 28.85 | 0.0177 | 717 |
| 14 | 22339422 | 104673942 | q11.2 | q32.33 | Gain | 82334521 | 13.74 | 0.00 | 13.74 | 0.0172 | 807 |
| 16 | 26570433 | 35271725 | p12.1 | p11.2 | Gain | 8701293 | 25.35 | 5.86 | 19.49 | 0.0211 | 86 |
| 16 | 46461309 | 74288643 | q11.2 | q22.3 | Gain | 27827335 | 28.23 | 8.15 | 20.08 | 0.0239 | 221 |
| 17 | 1 | 7296982 | p13.3 | p13.1 | AI | 7296982 | 23.25 | 4.32 | 18.93 | 0.0222 | 162 |
| 17 | 52573434 | 68833208 | q22 | q24.3 | Loss | 16259775 | 12.40 | 0.23 | 12.17 | 0.0285 | 198 |
| 20 | 0 | 7789904 | p13 | p12.3 | Gain | 7789905 | 52.58 | 30.39 | 22.18 | 0.0468 | 24 |

AI, Allelic imbalance

**Table S3. List of the identified somatic pathogenic single-nucleotide variants and Indels analysed by targeted NGS**

| Gene | Chrom | Position | CDS_Variant | Aminonacid_Variant | Effect |
|---|---|---|---|---|---|
| AKT1 | 14 | 105246551 | c.49G>A | p.Glu17Lys | Missense |
| AMER1 | X | 63411678 | c.1489C>T | p.Arg497* | Nonsense |
| AMER1 | X | 63412110 | c.1057C>T | p.Arg353* | Nonsense |
| APC | 5 | 112173917 | c.2626C>T | p.Arg876* | Nonsense |
| APC | 5 | 112173704 | c.2413C>T | p.Arg805* | Nonsense |
| APC | 5 | 112155002 | c.1275delA | p.Ala426fs | Indel_Frameshift |
| APC | 5 | 112178940 | c.7649A>G | p.Glu2550Gly | Missense |
| DLC1 | 8 | 13357402 | c.179C>T | p.Ser60Leu | Missense |
| APC | 5 | 112128191 | c.694C>T | p.Arg232* | Nonsense |
| APC | 5 | 112128191 | c.694C>T | p.Arg232* | Nonsense |
| APC | 5 | 112151261 | c.904C>T | p.Arg302* | Nonsense |
| APC | 5 | 112175576 | c.4285C>T | p.Gln1429* | Nonsense |
| APC | 5 | 112177379 | c.6088C>T | p.Leu2030Phe | Missense |
| APC | 5 | 112111373 | c.470G>A | p.Trp157* | Nonsense |
| APC | 5 | 112175162 | c.3871C>T | p.Gln1291* | Nonsense |
| APC | 5 | 112175466 | c.4175C>A | p.Ser1392* | Nonsense |
| APC | 5 | 112175639 | c.4348C>T | p.Arg1450* | Nonsense |
| APC | 5 | 112157682 | c.1402G>T | p.Glu468* | Nonsense |
| APC | 5 | 112175525 | c.4234G>T | p.Gly1412* | Nonsense |
| APC | 5 | 112175479 | c.4192_4193delAG | p.Arg1399fs | Indel_Frameshift |
| APC | 5 | 112174112 | c.2821G>T | p.Glu941* | Nonsense |
| APC | 5 | 112175903 | c.4612G>T | p.Glu1538* | Nonsense |
| APC | 5 | 112128143 | c.646C>T | p.Arg216* | Nonsense |
| APC | 5 | 112154943 | c.1214G>A | p.Arg405Gln | Missense |
| APC | 5 | 112175599 | c.4311_4312dupAA | p.Thr1438fs | Indel_Frameshift |
| APC | 5 | 112175639 | c.4348C>T | p.Arg1450* | Nonsense |
| APC | 5 | 112175328 | c.4037C>A | p.Ser1346* | Nonsense |
| APC | 5 | 112175273 | c.3982C>T | p.Gln1328* | Nonsense |
| ARID1A | 1 | 27101690 | c.4972C>T | p.Arg1658Trp | Missense |
| ARID1A | 1 | 27100877 | c.4159G>A | p.Glu1387Lys | Missense |
| ATM | 11 | 108121796 | c.1604C>T | p.Ser535Leu | Missense |
| ATM | 11 | 108121483 | c.1291G>A | p.Glu431Lys | Missense |
| ATM | 11 | 108160402 | c.4310G>T | p.Arg1437Ile | Missense |
| BRAF | 7 | 140481402 | c.1406G>C | p.Gly469Ala | Missense |
| BRAF | 7 | 140453155 | c.1780G>A | p.Asp594Asn | Missense |
| BRCA2 | 13 | 32893318 | c.172G>A | p.Glu58Lys | Missense |
| BRCA2 | 13 | 32944558 | c.8351G>A | p.Arg2784Gln | Missense |
| CDKN2A | 9 | 21971099 | c.425C>T | p.Pro142Leu | Missense |
| CSMD1 | 8 | 3205599 | c.3392G>A | p.Cys1131Tyr | Missense |
| CSMD1 | 8 | 3351211 | c.1385G>A | p.Arg462Gln | Missense |
| CSMD1 | 8 | 3351221 | c.1375G>A | p.Glu459Lys | Missense |
| CSMD1 | 8 | 2964134 | c.6868G>A | p.Gly2290Arg | Missense |
| CSMD1 | 8 | 3257027 | c.2294C>T | p.Ala765Val | Missense |
| CSMD1 | 8 | 2965294 | c.6784C>G | p.Pro2262Ala | Missense |
| CSMD1 | 8 | 2800063 | c.10469C>T | p.Ala3490Val | Missense |

| | | | | | |
|---|---|---|---|---|---|
| CSMD1 | 8 | 3267070 | c.1622G>A | | Missense |
| CTNNB1 | 3 | 41278084 | c.1960T>G | | Missense |
| CTNNB1 | 3 | 41277844 | c.1808T>G | p.Leu603Arg | Missense |
| CTNNB1 | 3 | 41274911 | c.1161T>A | p.Asn387Lys | Missense |
| FAT1 | 4 | 187540578 | c.7162C>T | p.Pro2388Ser | Missense |
| DAPK1 | 9 | 90261428 | c.1184T>G | p.Leu395Arg | Missense |
| DCC | 18 | 50866236 | c.1907G>A | p.Arg636Gln | Missense |
| APC | 5 | 112175828 | c.4537G>T | p.Glu1513* | Nonsense |
| APC | 5 | 112174481 | c.3190G>T | p.Glu1064* | Nonsense |
| APC | 5 | 112174094 | c.2804dupA | p.Tyr935fs | Indel_Frameshift |
| APC | 5 | 112175951 | c.4666dupA | p.Thr1556fs | Indel_Frameshift |
| EGFR | 7 | 55268909 | c.2975C>T | p.Pro992Leu | Missense |
| ERBB2 | 17 | 37881022 | c.2351G>A | p.Arg784His | Missense |
| FAT1 | 4 | 187540515 | c.7225G>T | p.Val2409Leu | Missense |
| FAT1 | 4 | 187540074 | c.7666C>T | p.Arg2556* | Nonsense |
| FAT1 | 4 | 187518871 | c.128C>T | p.Pro43Leu | Missense |
| FAT4 | 4 | 126371772 | c.9601C>T | p.Pro3201Ser | Missense |
| FAT4 | 4 | 126384750 | c.11827G>T | p.Glu3943* | Nonsense |
| FAT4 | 4 | 126242566 | c.5000T>C | p.Ile1667Thr | Missense |
| FAT4 | 4 | 126389866 | c.12099_12100insTG | p.Arg4034fs | Indel_Frameshift |
| FAT4 | 4 | 126238899 | c.1333G>A | p.Gly445Arg | Missense |
| FBXW7 | 4 | 153245380 | c.1811A>C | p.Lys604Thr | Missense |
| FBXW7 | 4 | 153258983 | c.832C>T | p.Arg278* | Nonsense |
| FLT3 | 13 | 28610098 | c.1392G>A | p.Trp464* | Nonsense |
| FLT3 | 13 | 28599027 | c.2261G>A | p.Gly754Glu | Missense |
| IRS2 | 13 | 110437803 | c.598G>A | p.Val200Met | Missense |
| IRS2 | 13 | 110435655 | c.2746C>A | p.Pro916Thr | Missense |
| KMT2C | 7 | 151842343 | c.14240G>A | p.Arg4747Gln | Missense |
| KMT2C | 7 | 151945007 | c.2512G>A | p.Gly838Ser | Missense |
| KMT2C | 7 | 151945007 | c.2512G>A | p.Gly838Ser | Missense |
| KMT2C | 7 | 151874739 | c.7799C>T | p.Pro2600Leu | Missense |
| KMT2C | 7 | 151843704 | c.14182G>A | p.Val4728Met | Missense |
| KMT2C | 7 | 151945007 | c.2512G>A | p.Gly838Ser | Missense |
| KMT2D | 12 | 49425232 | c.13256C>T | p.Pro4419Leu | Missense |
| KMT2D | 12 | 49422927 | c.14168C>A | p.Pro4723His | Missense |
| KMT2D | 12 | 49427702 | c.10786C>T | p.Arg3596Trp | Missense |
| KRAS | 12 | 25398281 | c.38G>A | p.Gly13Asp | Missense |
| KRAS | 12 | 25378647 | c.351A>C | p.Lys117Asn | Missense |
| KRAS | 12 | 25398284 | c.35G>A | p.Gly12Asp | Missense |
| KRAS | 12 | 25398262 | c.57G>T | p.Leu19Phe | Missense |
| KRAS | 12 | 25398284 | c.35G>A | p.Gly12Asp | Missense |
| KRAS | 12 | 25398284 | c.35G>A | p.Gly12Asp | Missense |
| KRAS | 12 | 25398284 | c.35G>A | p.Gly12Asp | Missense |
| KRAS | 12 | 25398284 | c.35G>A | p.Gly12Asp | Missense |
| KRAS | 12 | 25398284 | c.35G>A | p.Gly12Asp | Missense |
| KRAS | 12 | 25398281 | c.38G>A | p.Gly13Asp | Missense |
| KRAS | 12 | 25398285 | c.34G>T | p.Gly12Cys | Missense |
| KRAS | 12 | 25398284 | c.35G>A | p.Gly12Asp | Missense |
| KRAS | 12 | 25378562 | c.436G>C | p.Ala146Pro | Missense |

| | | | | | |
|---|---|---|---|---|---|
| KRAS | 12 | 25398284 | c.35G>T | p.Gly12Val | Missense |
| KRAS | 12 | 25398284 | c.35G>A | p.Gly12Asp | Missense |
| KRAS | 12 | 25398281 | c.38G>A | p.Gly13Asp | Missense |
| KRAS | 12 | 25398285 | c.34G>T | p.Gly12Cys | Missense |
| KRAS | 12 | 25398281 | c.38G>A | p.Gly13Asp | Missense |
| KRAS | 12 | 25398284 | c.35G>T | p.Gly12Val | Missense |
| KRAS | 12 | 25378562 | c.436G>A | p.Ala146Thr | Missense |
| KRAS | 12 | 25398284 | c.35G>A | p.Gly12Asp | Missense |
| KRAS | 12 | 25398284 | c.35G>A | p.Gly12Asp | Missense |
| LRP1B | 2 | 141115581 | c.11362G>A | p.Gly3788Arg | Missense |
| LRP1B | 2 | 141819784 | c.1072G>A | p.Gly358Arg | Missense |
| LRP1B | 2 | 141272273 | c.8218G>A | p.Asp2740Asn | Missense |
| LRP1B | 2 | 141215043 | c.9803G>A | p.Arg3268Lys | Missense |
| LRP1B | 2 | 141709502 | c.2894delC | p.Pro965fs | Indel_Frameshift |
| LRP1B | 2 | 141762970 | c.2437C>A | p.Pro813Thr | Missense |
| LRP1B | 2 | 141762918 | c.2489G>A | p.Gly830Glu | Missense |
| LRP1B | 2 | 141460072 | c.6074G>C | p.Arg2025Pro | Missense |
| LRP1B | 2 | 141812703 | c.1534G>A | p.Asp512Asn | Missense |
| LRP1B | 2 | 141298575 | c.7480G>T | p.Asp2494Tyr | Missense |
| MAP2K7 | 19 | 7974652 | c.137C>T | p.Pro46Leu | Missense |
| MET | 7 | 116435814 | c.3958C>T | p.Leu1320Phe | Missense |
| MLH1 | 3 | 37092065 | c.2192C>T | p.Pro731Leu | Missense |
| MLH1 | 3 | 37061871 | c.955G>A | p.Glu319Lys | Missense |
| MSH6 | 2 | 48027661 | c.2539G>A | p.Glu847Lys | Missense |
| MSH6 | 2 | 48030612 | c.3226C>T | p.Arg1076Cys | Missense |
| NOTCH1 | 9 | 139405141 | c.2704C>T | p.Arg902Cys | Missense |
| NOTCH1 | 9 | 139404225 | c.2929G>A | p.Gly977Arg | Missense |
| NOTCH1 | 9 | 139401210 | c.3859C>T | p.Arg1287Cys | Missense |
| NOTCH1 | 9 | 139412240 | c.1405G>T | p.Asp469Tyr | Missense |
| NOTCH1 | 9 | 139409816 | c.1940G>A | p.Ser647Asn | Missense |
| NOTCH1 | 9 | 139404272 | c.2882C>T | p.Thr961Met | Missense |
| NOTCH1 | 9 | 139402531 | c.3386A>T | p.His1129Leu | Missense |
| NOTCH1 | 9 | 139401871 | c.3529G>A | p.Gly1177Arg | Missense |
| NOTCH1 | 9 | 139391982 | c.6209G>A | p.Arg2070Gln | Missense |
| PCLO | 7 | 82763648 | c.3218C>T | p.Pro1073Leu | Missense |
| PCLO | 7 | 82430843 | c.14998G>A | p.Asp5000Asn | Missense |
| PCLO | 7 | 82430851 | c.14990G>A | p.Gly4997Glu | Missense |
| PCLO | 7 | 82763597 | c.3269G>A | p.Cys1090Tyr | Missense |
| PIK3CA | 3 | 178936082 | c.1624G>A | p.Glu542Lys | Missense |
| PIK3CA | 3 | 178936082 | c.1624G>A | p.Glu542Lys | Missense |
| PIK3CA | 3 | 178936091 | c.1633G>A | p.Glu545Lys | Missense |
| PIK3CA | 3 | 178952013 | c.3068G>A | p.Arg1023Gln | Missense |
| POLE | 12 | 133252391 | c.1036A>T | p.Arg346Trp | Missense |
| PTEN | 10 | 89692935 | c.419T>C | p.Leu140Ser | Missense |
| RNF43 | 17 | 56435476 | c.1661G>A | p.Arg554Gln | Missense |
| ROBO1 | 3 | 78737847 | c.1121C>T | p.Ala374Val | Missense |
| ROBO1 | 3 | 78737868 | c.1100C>A | p.Thr367Asn | Missense |
| ROBO1 | 3 | 78649393 | c.4811C>T | p.Ser1604Leu | Missense |
| SETD2 | 3 | 47147573 | c.4753G>A | p.Asp1585Asn | Missense |

| | | | | | |
|---|---|---|---|---|---|
| SMAD2 | 18 | 45371786 | c.1205T>C | p.Phe402Ser | Missense |
| SMAD2 | 18 | 45368211 | c.1391C>G | p.Ser464* | Nonsense |
| SMAD4 | 18 | 48604747 | c.1569C>A | p.Cys523* | Nonsense |
| SMAD4 | 18 | 48591919 | c.1082G>A | p.Arg361His | Missense |
| SMAD4 | 18 | 48604788 | c.1610A>G | p.Asp537Gly | Missense |
| SOX9 | 17 | 70120093 | c.1100delC | p.Pro367fs | Indel_Frameshift |
| SOX9 | 17 | 70118934 | c.506A>C | p.His169Pro | Missense |
| SOX9 | 17 | 70119932 | c.934C>T | p.Gln312* | Nonsense |
| SOX9 | 17 | 70120178 | c.1180C>T | p.Arg394* | Nonsense |
| SOX9 | 17 | 70120139 | c.1143_1144dupGC | p.Leu382fs | Indel_Frameshift |
| SOX9 | 17 | 70117614 | c.82G>T | p.Glu28* | Nonsense |
| SOX9 | 17 | 70119814 | c.818_819dupTG | p.Asp274fs | Indel_Frameshift |
| SOX9 | 17 | 70120040 | c.1042C>T | p.Gln348* | Nonsense |
| SOX9 | 17 | 70120041 | c.1049_1058delCACAGGCCCC | p.Pro350fs | Indel_Frameshift |
| SYNE1 | 6 | 152629725 | c.17245C>T | p.His5749Tyr | Missense |
| SYNE1 | 6 | 152532699 | c.22519C>T | p.Arg7507Cys | Missense |
| SYNE1 | 6 | 152510440 | c.23248C>T | p.Leu7750Phe | Missense |
| SYNE1 | 6 | 152563412 | c.19856C>G | p.Ser6619Cys | Missense |
| SYNE1 | 6 | 152615120 | c.17825G>A | p.Arg5942His | Missense |
| SYNE1 | 6 | 152644776 | c.15754G>A | p.Glu5252Lys | Missense |
| SYNE1 | 6 | 152462425 | c.25159G>A | p.Val8387Met | Missense |
| SYNE1 | 6 | 152751744 | c.4562G>A | p.Arg1521Gln | Missense |
| TP53 | 17 | 7576569 | c.1009C>T | p.Arg337Cys | Missense |
| TP53 | 17 | 7574003 | c.1024C>T | p.Arg342* | Nonsense |
| TP53 | 17 | 7579406 | c.281C>A | p.Ser94* | Nonsense |
| TP53 | 17 | 7578478 | c.452C>A | p.Pro151His | Missense |
| TP53 | 17 | 7578475 | c.455C>T | p.Pro152Leu | Missense |
| TP53 | 17 | 7578452 | c.472_477delCGCGCC | p.Arg158_Ala159del | Indel_Inframe |
| TP53 | 17 | 7578407 | c.523C>G | p.Arg175Gly | Missense |
| TP53 | 17 | 7578406 | c.524G>A | p.Arg175His | Missense |
| TP53 | 17 | 7578406 | c.524G>A | p.Arg175His | Missense |
| TP53 | 17 | 7578406 | c.524G>A | p.Arg175His | Missense |
| TP53 | 17 | 7578395 | c.535C>A | p.His179Asn | Missense |
| TP53 | 17 | 7578236 | c.613T>G | p.Tyr205Asp | Missense |
| TP53 | 17 | 7578211 | c.638G>T | p.Arg213Leu | Missense |
| TP53 | 17 | 7578208 | c.641A>G | p.His214Arg | Missense |
| TP53 | 17 | 7577548 | c.733G>A | p.Gly245Ser | Missense |
| TP53 | 17 | 7577548 | c.733G>A | p.Gly245Ser | Missense |
| TP53 | 17 | 7577547 | c.734G>A | p.Gly245Asp | Missense |
| TP53 | 17 | 7577539 | c.742C>T | p.Arg248Trp | Missense |
| TP53 | 17 | 7579715 | c.80delC | p.Pro27fs | Indel_Frameshift |
| TP53 | 17 | 7577121 | c.817C>T | p.Arg273Cys | Missense |
| TP53 | 17 | 7577121 | c.817C>T | p.Arg273Cys | Missense |
| TP53 | 17 | 7577106 | c.832C>A | p.Pro278Thr | Missense |
| TP53 | 17 | 7577094 | c.844C>T | p.Arg282Trp | Missense |
| TP53 | 17 | 7577022 | c.916C>T | p.Arg306* | Nonsense |
| TCF7L2 | 10 | 114912181 | c.1251G>T | p.Trp417Cys | Missense |
| DLC1 | 8 | 12958102 | c.1744C>A | p.Leu582Met | Missense |
| ATM | 11 | 108098418 | c.67C>T | p.Arg23* | Nonsense |

#### Supplementary Figure S1

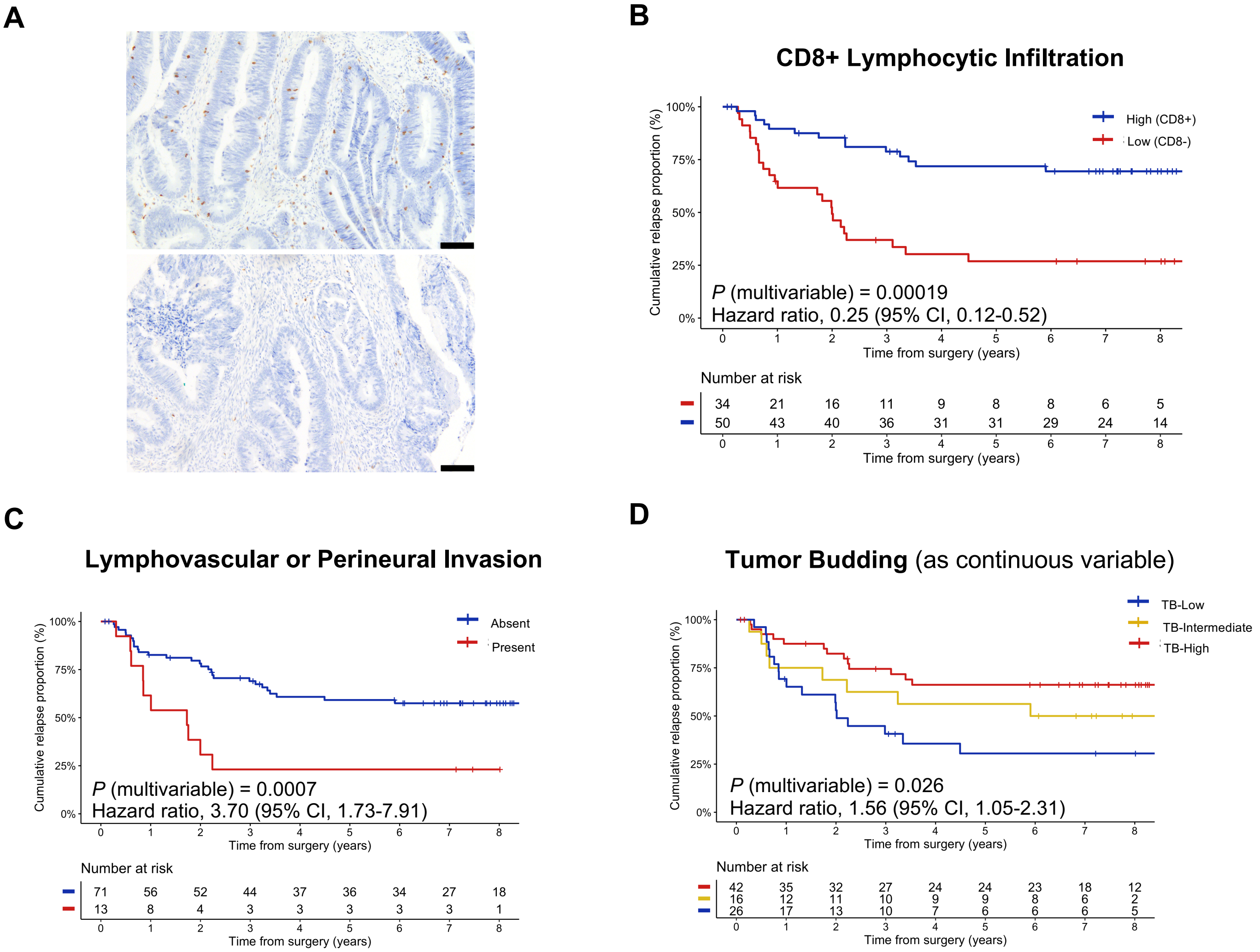

#### Supplementary Figure S2

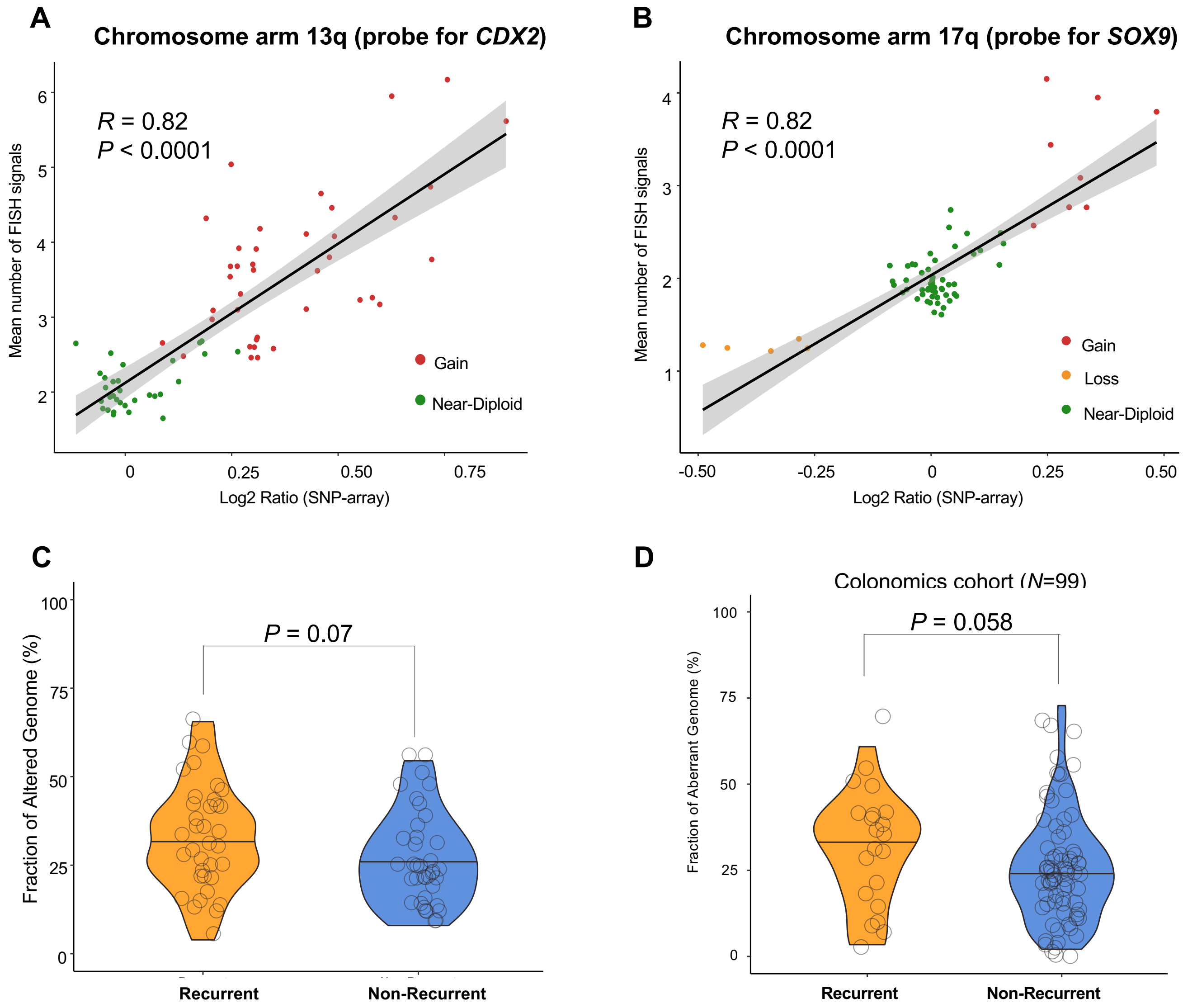

### Supplementary Figure S3

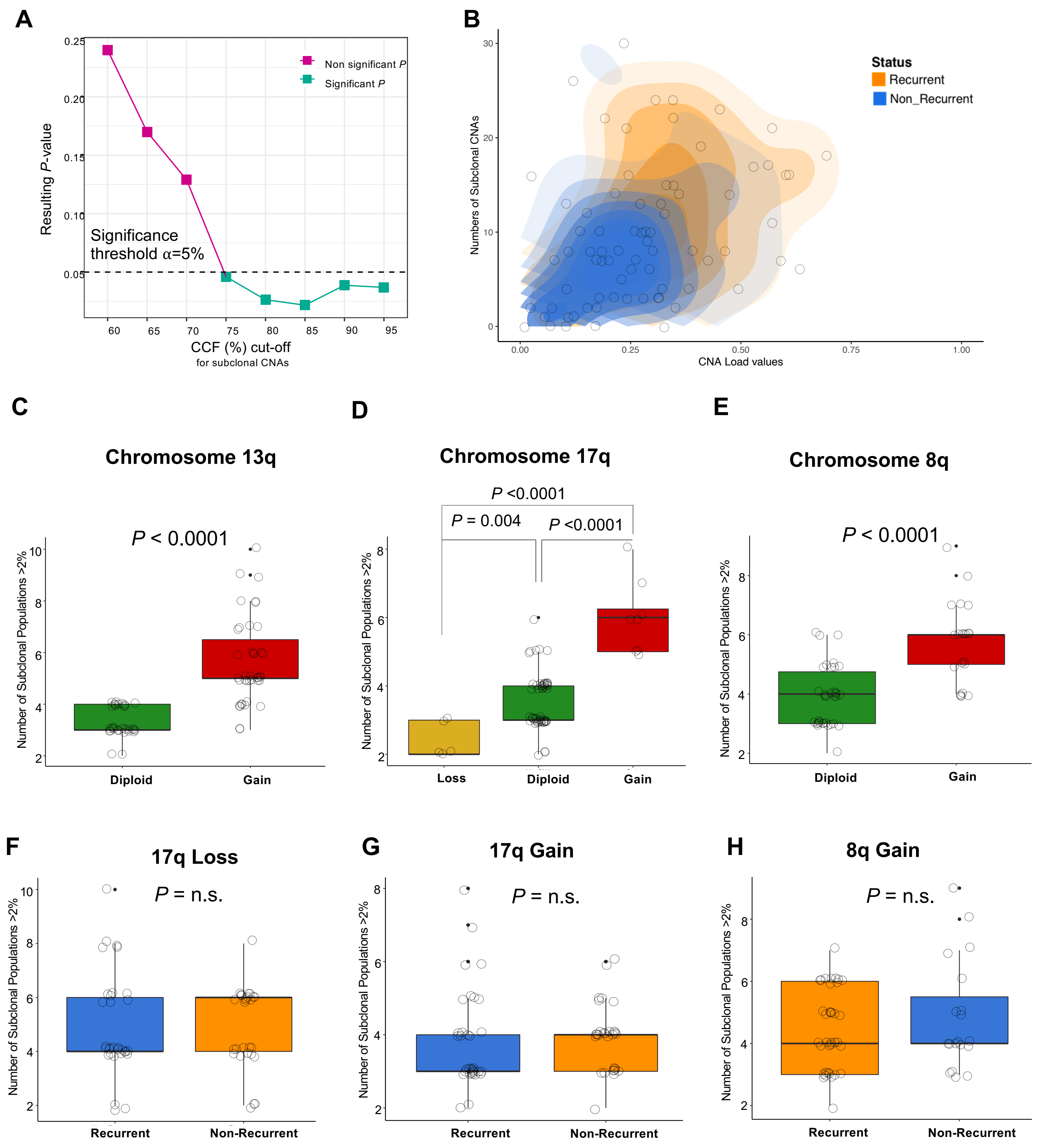

### Supplementary Figure S4

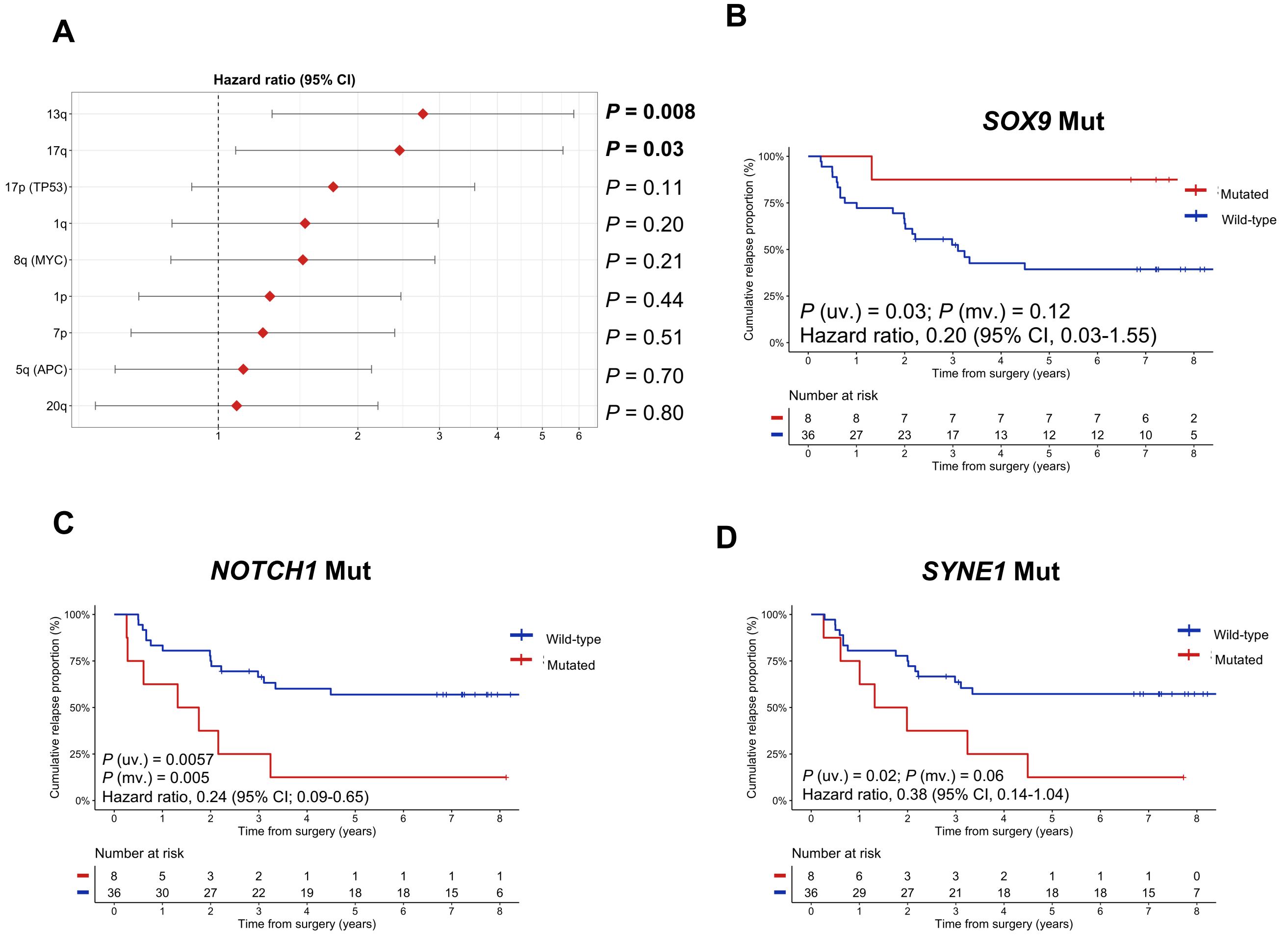

### Supplementary Figure S5

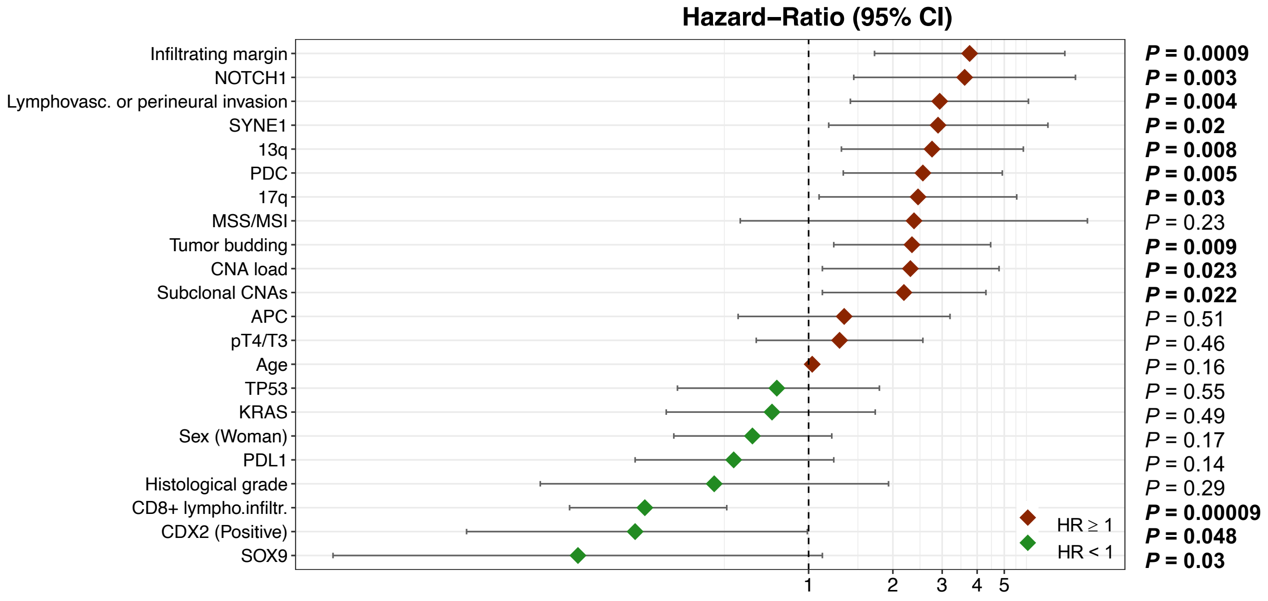

**Supplementary Figure S6**

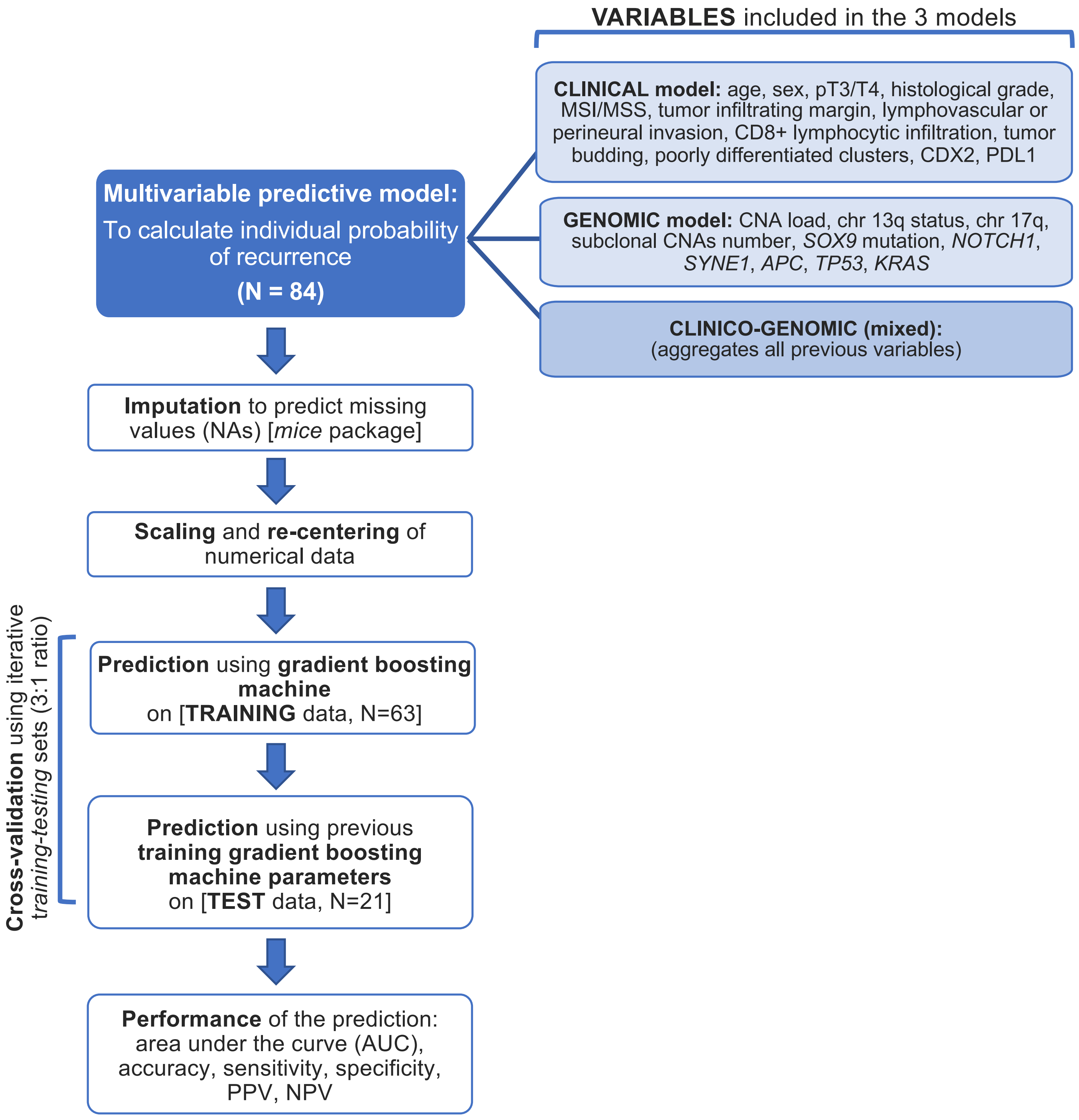
